# Loss of microbial diversity and body site heterogeneity in individuals with Hidradenitis Suppurativa

**DOI:** 10.1101/612457

**Authors:** Andrea M. Schneider, Lauren C. Cook, Xiang Zhan, Kalins Banerjee, Zhaoyuan Cong, Yuka Imamura-Kawasawa, Samantha L. Gettle, Amy L. Longenecker, Joslyn S. Kirby, Amanda M. Nelson

## Abstract

Hidradenitis Suppurativa (HS) is a chronic, scarring, inflammatory skin disease affecting hair follicles in axillae, inguinal, and anogenital regions. Dysbiosis in HS patients compared to healthy subjects is documented. However, whether dysbiosis is specific to particular body sites or skin niches is unknown. We investigated the follicular and skin surface microbiome of the axilla and groin of HS patients (n=11) and healthy individuals (n=10) using 16S rRNA gene sequencing (V3-V4). We sampled non-lesional (HSN) and lesional skin (HSL) of HS patients. β-diversity was significantly decreased (p<0.05) in HSN and HSL skin compared to normal skin with loss of body site and skin niche heterogeneity in HS samples. The relative bacterial abundance of specific microbes was also significantly different between normal and HSN (15 genera) or HSL (21 genera) skin. Smoking and alcohol use influenced the β-diversity (p<0.08) in HS skin. We investigated metabolic profiles of bacterial communities in HS and normal skin using a computational approach. *Metabolism, Genetic Information Processing*, and *Environmental Information Processing* were significantly different between normal and HS samples. Altered metabolic pathways associated with dysbiosis of HS skin suggest mechanisms underlying the disease pathology and information about treatment with drugs targeting those pathways.

## INTRODUCTION

Hidradenitis Suppurativa (HS) is a chronic scarring inflammatory skin disease affecting the pilosebaceous units (PSU) of the axilla, inframammary folds, groin, and buttocks. Clinical features include painful nodules, abscesses, draining sinus tracts, and fistulas. Antibiotics are commonly prescribed to treat HS, however, lesions are often sterile. With the increased prevalence of antibiotic resistance, understanding the role of bacteria in HS is critical.

The skin microbiome varies by individual, body site, age, and disease status (Grice et al. 2009; Kong et al. 2012; Oh et al. 2014). Dysbiosis, or microbial imbalance, of the skin microbiome occurs in atopic dermatitis, epidermolysis bullosa, chronic non-healing skin wounds, and others (Fuentes et al. 2018; Kalan et al. 2016; Kong et al. 2012).

Dysbiosis has also been reported in HS (Guet-Revillet et al. 2017; Nikolakis et al. 2017; Ring et al. 2017b; Ring et al. 2017a). Collectively, the HS microbiome has been studied by invasive and non-invasive sampling of non-lesional and lesional skin using culture and next generation sequencing analyses. The only consistent finding between these studies was that bacterial dysbiosis exists in HS skin compared to normal skin. However, where in the skin the dysbiosis is occurring and whether or not this dysbiosis alters microbiome functions remains unexplored.

In this study, we investigated whether dysbiosis is specific to a particular body site or skin niche and applied a computational approach to understand the biological impact of dysbiotic communities in HS patients. We sampled both the follicular and skin surface microbiome of the axilla and groin of HS patients (n=11) and healthy individuals (n=10) followed by 16S rRNA gene sequencing (V3-V4). This study reveals that loss of bacterial diversity in HS patients extends to both skin niches. Alterations in the metabolic profile due to dysbiosis may contribute to HS symptoms and pathology.

## RESULTS

### Study subjects and sequencing methods

Ten normal subjects and 11 subjects with HS were enrolled; including males and females, ages 18-60 years after providing informed written consent under an IRB approved protocol. The diagnosis of HS was confirmed by a dermatologist. For subjects with HS, the Hurley Stage, disease duration, and family history of HS and other inflammatory conditions were documented. Use of medications, alcohol, hygiene products and smoking history were documented for all subjects **(Table S1)**.

Cyanoacrylate follicular biopsy (glue) and swab samples were collected from the axilla vault and inguinal crease (groin) for each individual. Four samples were collected from each normal individual, while both non-lesional (HSN) and lesional (HSL) skin was sampled in each HS patient (8 samples) (**Figure 1a**). HSN skin appeared clinically normal and was ∼5 cm away from an active lesion; lesional skin was defined by an inflammatory nodule. We specifically chose this sampling distance to keep within the same anatomical body area (ie: axilla vault). A total of 128 patient samples, four negative sampling controls, and one positive mock community control (MCC) (**Figure S1**) were collected.

**Figure 1:**
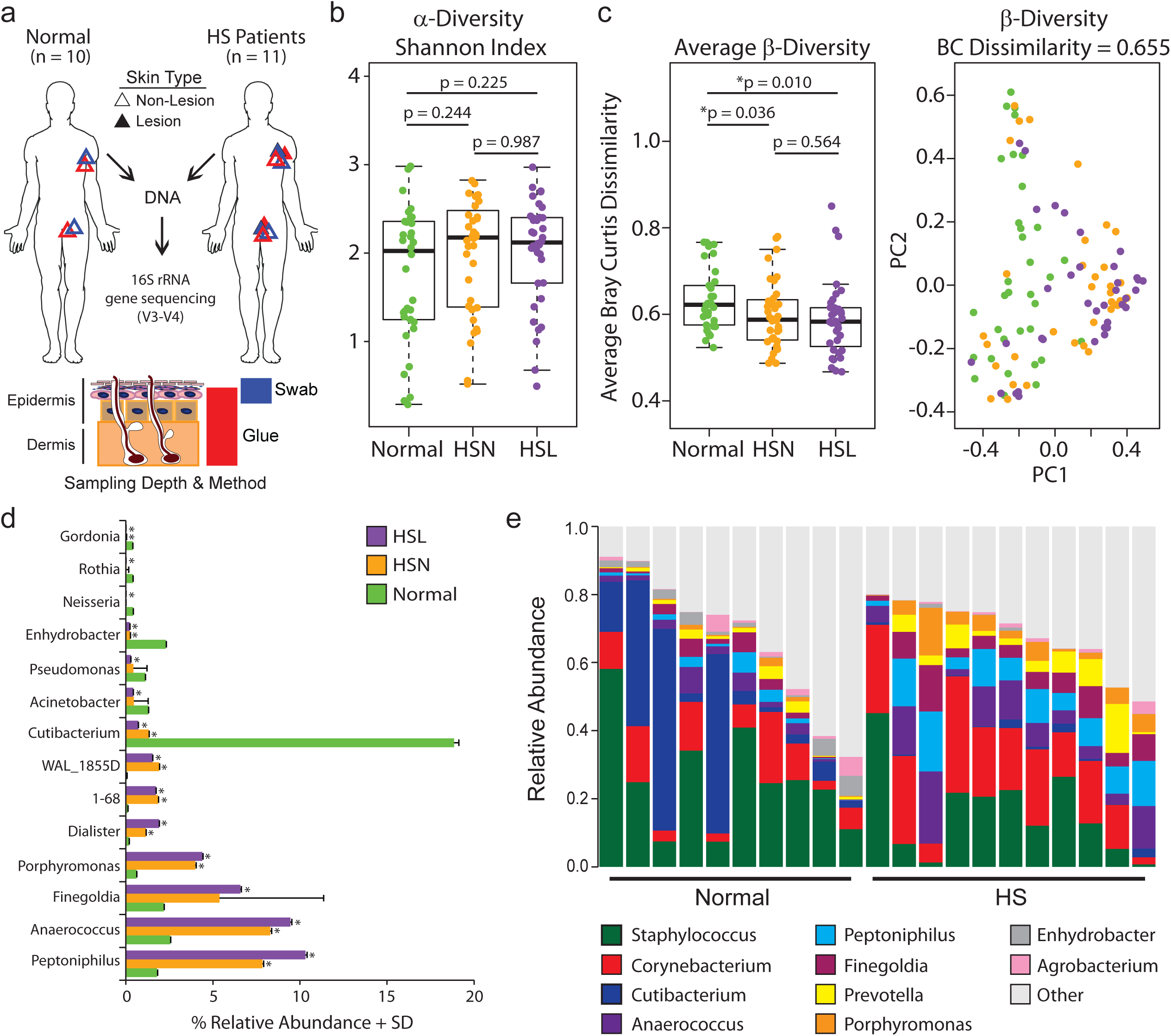
Bacterial β-diversity is significantly decreased on HS compared to normal skin. **a)** Schematic of experimental design **b)** Shannon diversity index (α-diversity) for normal (n=34), HSN (n=36), and HSL (n=36) skin samples; Wilcoxon rank sum test, p < 0.05 considered significant. HSN = HS Non-lesional, HSL = HS Lesional. **c)** Bray-Curtis dissimilarity index for β-diversity comparison, Wilcoxon rank sum test, p < 0.05 significant. The Bray-Curtis dissimilarity index is a scale from 0 to 1: 0 indicates identical communities and 1 indicates completely dissimilar communities. **d)** Top 20 significantly different genera between normal, HSL, and HSN skin. Mean relative abundance ± SD: * significantly different from normal. **e)** Relative bacterial abundance of the top 10 genera of each subject.

16S rRNA gene sequencing V1-V3 is preferred in skin microbiome studies, while V4 is more common for gut microbiome research (Meisel et al. 2016). We chose the V3-V4 region (**Table S2; Supplemental Methods M3**) in an effort to maximize identification of both skin and gut communities, as common gut microbiota have been reported within HS lesions (Ring et al. 2017b). Only samples with a minimum of 75,000 sequences were included in subsequent analyses (n = 117) (**Table S3**).

In total, we identified 994 operational taxonomic units (OTUs) following the convention of 97% similarity for OTU assignment. OTUs that did not reach an abundance of 0.5% or higher in at least one sample were excluded from further analysis as these were considered potential sequencing errors (Li 2015). After filtering, 272 OTUs became the focus of downstream statistical analysis, belonging to 146 distinct genera and 1 unclassified group. When body site or skin niche were compared, samples were paired by body site, skin niche, and non-lesional or lesional skin to account for intrapersonal variability.

### Bacterial β-diversity is significantly decreased on HS compared to normal skin

First, we compared bacterial composition between normal, HSN, and HSL regardless of sampling method and body site. There was no significant difference in α-diversity (Shannon diversity index) between normal, HSN, and HSL skin (**Figure 1b**). However, a significant loss of β-diversity (Bray-Curtis dissimilarity index) was observed in both HSN and HSL compared to normal skin (**Figure 1c**). Overall differences in diversity of the bacterial communities were assessed with the Microbiome Regression Kernel Association Test (MiRKAT) (Zhao et al. 2015). The community level composition in normal skin was distinctly different from the composition of both HSN (p=0.036) and HSL (p=0.010) skin (**Figure 1c**).

We investigated community level diversity between HSN and HSL skin. No statistical difference in bacterial composition between HSN and HSL skin (p=0.4404) was found, in contrast to prior reports (Guet-Revillet et al. 2017; Ring et al. 2017b). When comparing individual taxa between HSN and HSL skin for each individual, no single genus was significantly different under a family-wise error rate (FWER) of 0.05, indicating homogeneous bacterial compositions. Overall, the bacterial communities between HSN and HSL samples were highly similar with a loss of β-diversity in HS skin compared to normal skin.

The relative abundance of individual taxa were compared between HS and normal skin. Fifteen (9 of the top 20) genera were significantly different between normal and HSN samples and 21 (14 of the top 20) genera were significantly different between normal and HSL samples (**Figure 1d, Table S4, S5**). With the high degree of similarity between HSN and HSL skin, we averaged these samples together for each HS patient and compared the top 10 most abundant genera between HS skin and healthy skin (**Figure 1e**). *Staphylococcus* and *Corynebacterium* comprised ∼35% of the bacteria in both normal and HS skin; however, *Corynebacterium* was ∼2 fold more abundant in HS skin. Like previous studies, *Cutibacterium* (formally *Propionibacterium* (Scholz and Kilian 2016)) was more abundant in normal skin (18.8% vs 1%, p<0.05), and *Peptoniphilus* and *Porphyromonas* were more abundant in HS skin (Ring et al. 2017b) (**Figure 1e, Table S4, S5**). Interestingly, *Arcanobacterium* was detected in samples from 9 of 11 HS patients (mean relative abundance 0.07%) but not detected in any sample from normal subjects. Overall, the HS disease state affects the relative abundance of a number of skin commensals and opportunistic pathogens.

### Body site and skin niche heterogeneity is lost in HS skin compared to normal skin

Skin microbiota vary by anatomical body site in healthy individuals with the microenvironment influencing bacterial colonization; skin sites with similar moisture, lipids, and pH have a more similar composition than disparate sites (Grice et al. 2008). Moreover, the skin’s surface and follicular microbiomes are not 100% identical (Grice et al. 2008; Hall et al. 2018). Because HS is a disease of the PSU that commonly affects the axilla and groin, we investigated whether body site or skin niche are different in HS compared to normal subjects. For these analyses, we stratified our data by body site or skin niche and included only ‘paired’ samples to account for intrapersonal variability (**Figures 2-3, S2-3**). Within HS skin, whether HSN or HSL, we observed no difference in α- or β-diversity when comparing between axilla vs. groin and skin surface vs. follicle, highlighting the relative homogeneous bacterial populations at both anatomical body locations and within skin niches. However, in healthy individuals, there are differences between the axilla and groin as well as between skin’s surface and follicle. In general, α-diversity is increased in the groin compared to the axilla (**Figure 2**). Moreover, the skin’s surface is more diverse (α-diversity) than the follicle, especially within the axilla (**Figure 3**). These data highlight the importance of both sampling method and body site sampling to the outcome of microbiome studies. Taken together, these data demonstrate that there is a loss of heterogeneity between body sites and skin niches in HS patients compared to healthy controls.

**Figure 2:**
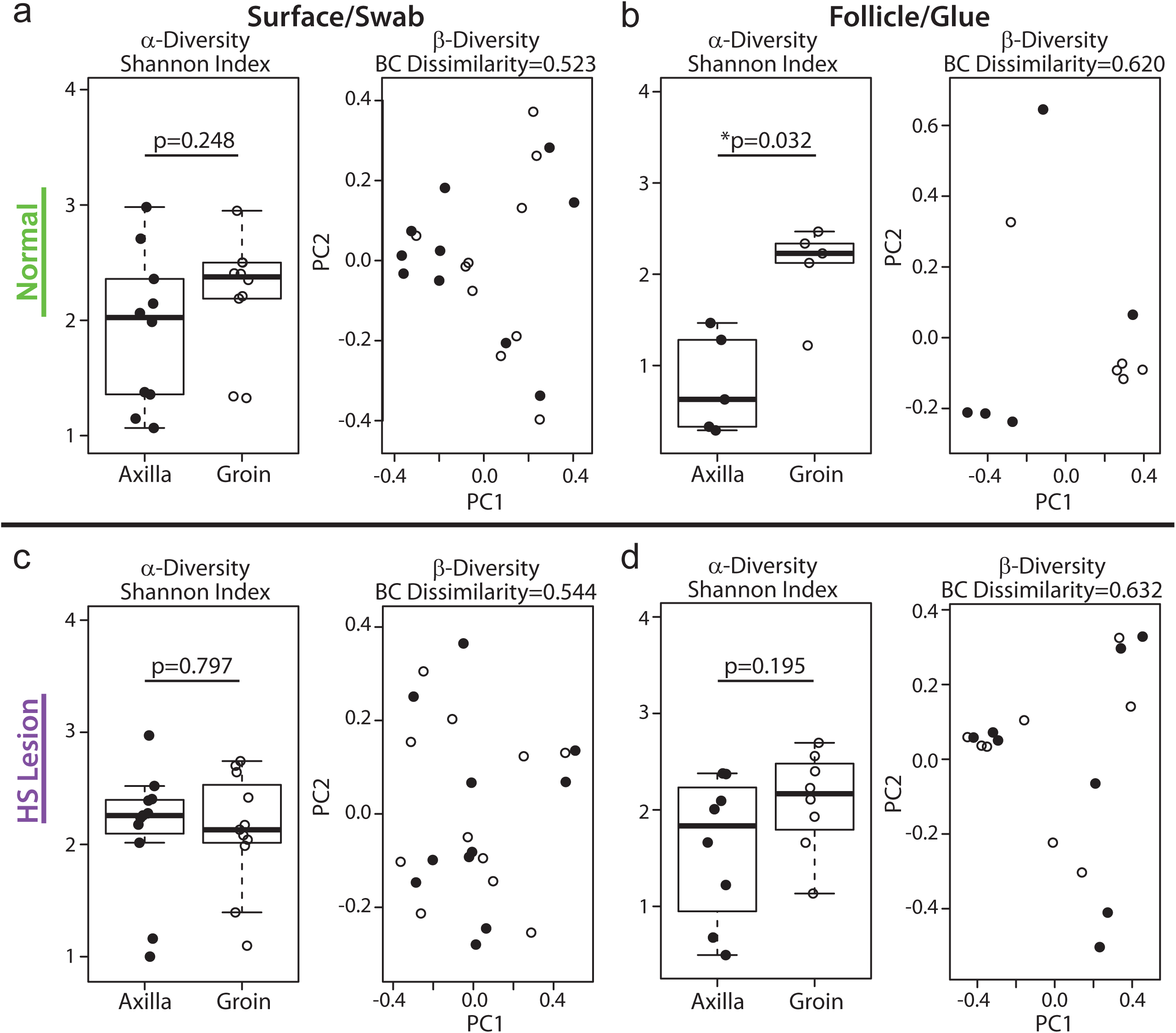
Body site heterogeneity is lost in HS compared to normal skin. **a-d)** Shannon diversity index (α-diversity) and Bray-Curtis dissimilarity index (β-diversity) comparison between axilla and groin stratified by normal and HSL skin and sampling method as indicated. The Wilcoxon rank sum test was applied to determine statistical differences in Shannon index between groups with p < 0.05 considered significant. **a)** Normal swab, n=10 from each body site, **b)** normal glue, n=5 from each body site, **c)** HS lesional swab, n=11 from each body site, and d) HS lesional glue, n=8 from each body site.

**Figure 3:**
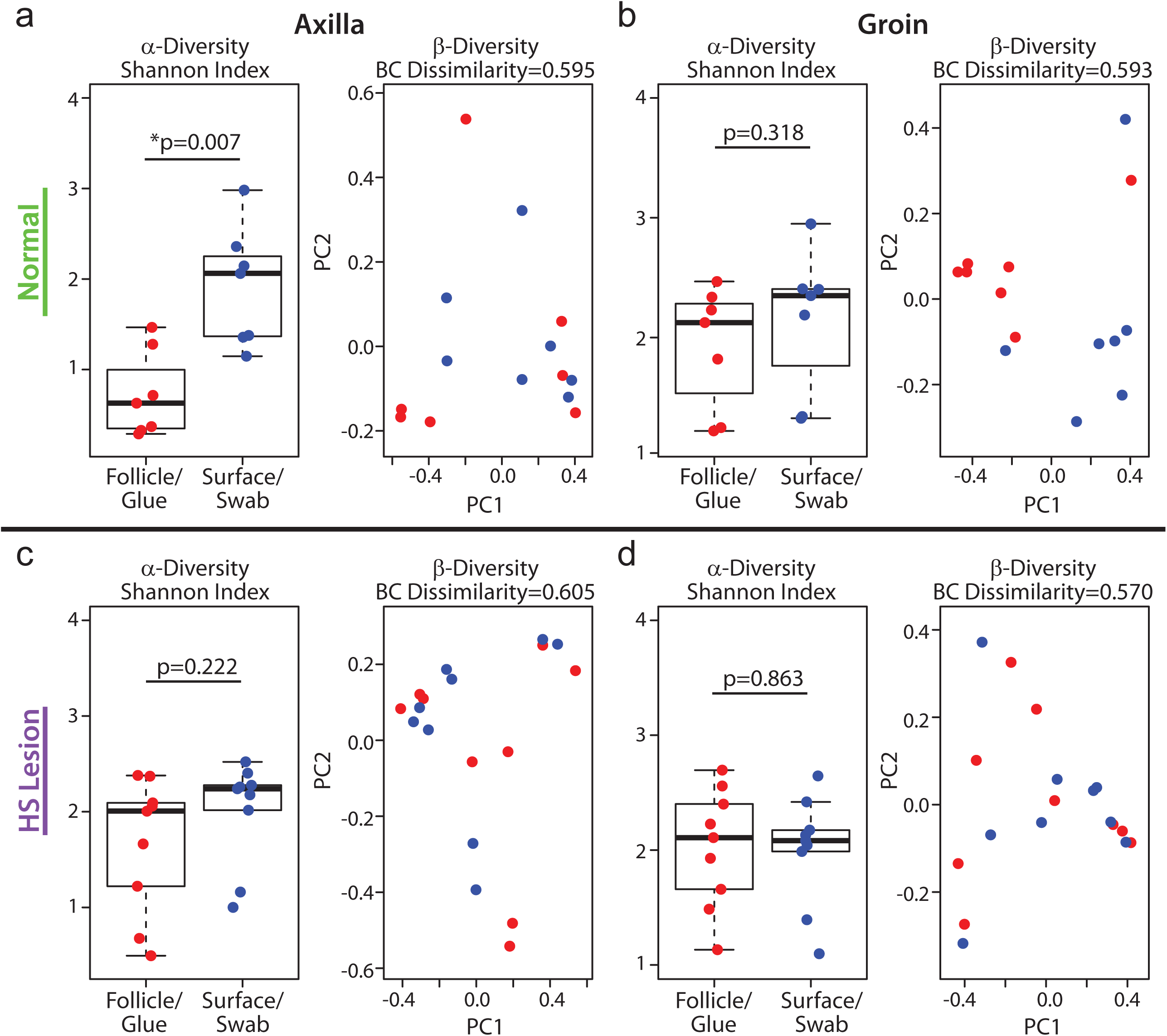
The follicle and surface microbiome signatures are different in normal skin, but not in HS skin. **a-d)** Shannon diversity index (α-diversity) and Bray-Curtis dissimilarity index (β-diversity) comparison between sampling methods (glue vs. swab) stratified by skin type and body site as indicated. The Wilcoxon rank sum test was applied to determine statistical differences in Shannon index between groups, with p < 0.05 considered significant. **a)** Normal axilla, n=7 collected by each method, **b)** normal groin, n=7 collected by each method, **c)** HS lesion axilla, n=9 collected by each method, and **d)** HS lesion groin, n=9 collected by each method.

### Individual lifestyle factors are associated with altered bacterial diversity in HS skin

We investigated correlations between bacterial diversity and subject demographics (**Table S1**). For each individual, we calculated the average relative abundance of each bacterial genus, including all samples regardless of location or sampling method. We used the MiRKAT association test (β-diversity) to investigate associations with sex, age, race, and use of hygiene, alcohol, or tobacco products (**Figure 4a**). This study was not sufficiently powered to detect significant changes associated with sex, race, or medication use (**Table S1**).

**Figure 4:**
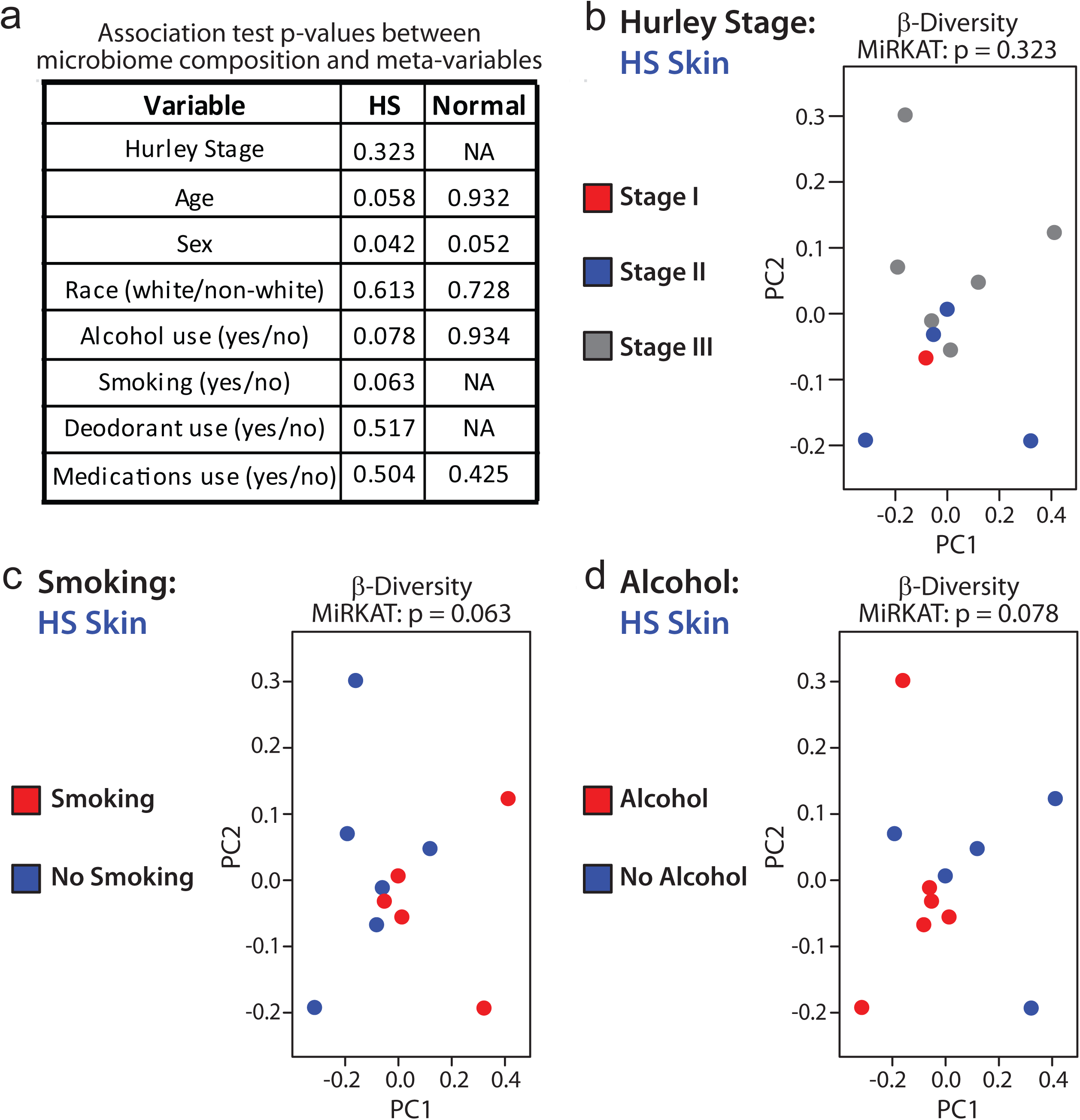
Individual lifestyle factors are associated with altered bacterial diversity in HS skin. **a)** MiRKAT analyses were applied to determine β-diversity differences (p-values) between normal subjects or HS patients associated with each variable as listed in the table. **b-d)** Bray-Curtis dissimilarity index (β-diversity) comparison in HS patients by Hurley stage, stage I (n=1, red), stage II (n=4, blue), stage III (n=6, grey) (**b**) smoking (n=5, red) vs. non-smoking (n=6, blue) **(c)** and between alcohol (n=6, red) vs. no alcohol (n=5, blue) **(d)**.

HS disease severity is classified by Hurley Stage (I, II, III). Previous studies demonstrated increased bacterial diversity with increasing Hurley Stage (Guet-Revillet et al. 2017; Nikolakis et al. 2017), however, due to limited sample size, no significant difference in β-diversity was found between disease stage in our study (**Figure 4a-b, Table S6**). The relative abundance of each genus was similar between Hurley Stage II and III except for *Prevotella* and *Atopobium,* which were significantly more abundant in stage III compared to stage II (**Table S7**).

Lifestyle choices may impact the microbiome. Both tobacco and alcohol use are linked to altered gut microbiomes (Capurso and Lahner 2017). However, the impact on the skin microbiome is not clear. Tobacco use is common in HS patients and is associated with increased disease severity (Akdogan et al. 2018; Sartorius et al. 2009). The number of HS patients who smoked (5 of 11) significantly differed from normal subjects (0 of 10) (**Table S6**). In HS patients, a trend in altered β-diversity (p = 0.063) between smoking and non-smoking was observed (**Figure 4c**). Alcohol use is also common among HS patients (Garg et al. 2018). Alcohol use also influenced bacterial diversity (p=0.078) between HS patients who were alcohol users vs. those who were not (6 and 5 patients, respectively) (**Figure 4d**). Together, in our HS patient population, these results suggest that lifestyle factors can impact skin microbiome diversity.

### KEGG Ontology indicates functional differences between microbiota of HS and normal skin

Like previous studies, our data illustrated decreased β-diversity between normal and HS skin. However, the biological impact of this altered diversity remains unexplored. Using a computational approach with PICRUSt, we investigated the metabolic profiles of HS and normal bacterial communities (Langille et al. 2013). We used KEGG pathway analysis to identify significant functional differences between the microbiota of normal, HSN, and HSL skin (**Figure 5a, Supplemental Methods M4**). Several KEGG Orthology groups were different between normal, HSN, and HSL samples: Level 1 (n=3), Level 2 (n=15), and Level 3 (n=52) (**Table S8**). At the highest categorical level, *Metabolism, Genetic Information Processing*, and *Environmental Information Processing* were significantly different between normal and HSN or HSL skin (**Table S8**). Several *Metabolism* pathways were enriched in normal compared to HS skin including: *Pyruvate, Arginine and proline; Propanoate; Retinol; Valine, leucine, and isoleucine biosynthesis;* and *Valine, leucine, and isoleucine degradation. Purine and pyrimidine metabolism* were significantly enriched in both HSN and HSL compared to normal skin (**Figure 5b**). Within *Genetic Information Processing*, all KEGG Ortholog terms were significantly enriched in HSN and HSL compared to the normal skin (**Figure 5c**); this combined with increased *purine and pyrimidine metabolism* suggests increased microbial turnover in HS compared to normal skin.

**Figure 5:**
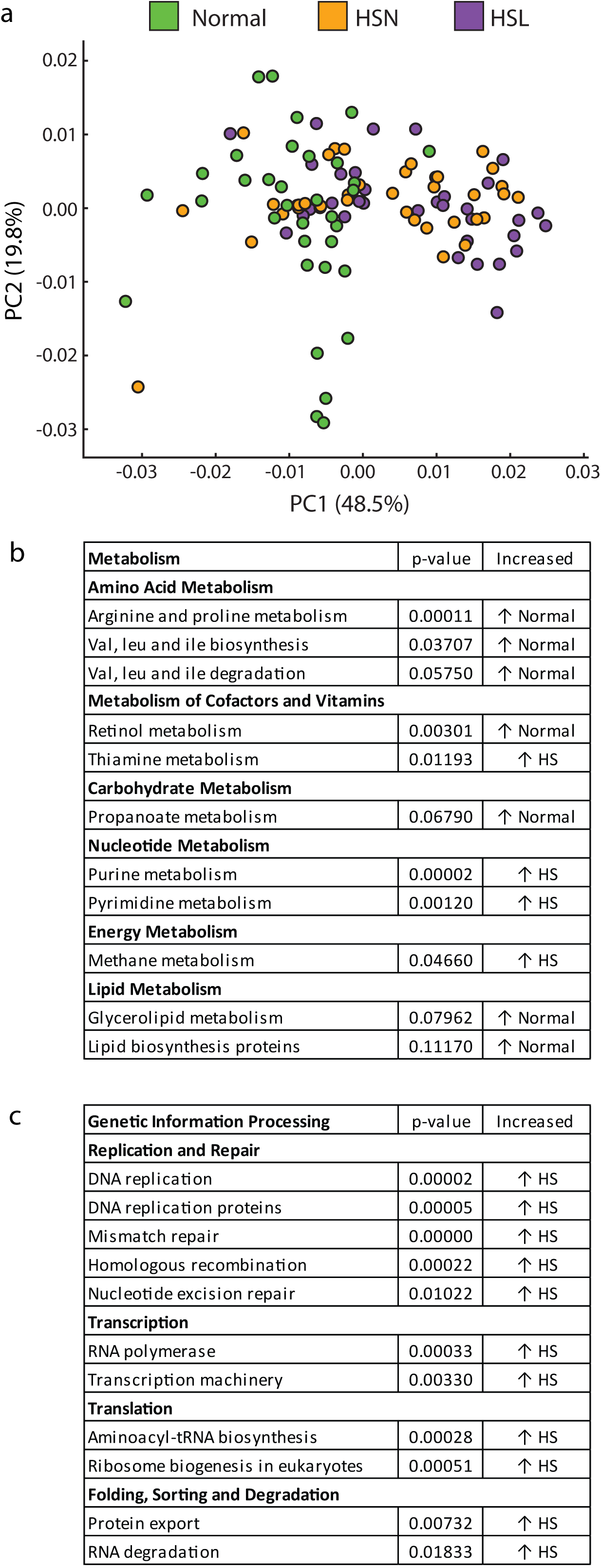
KEGG Ontology indicates functional differences between microbiota of HS and normal skin. **a)** PCA plot of the overall functional differences of Level 3 KEGG pathways between normal (n=34), HSN (n=36), and HSL (n=36) skin. Multiple group statistical analysis was performed in STAMP (Kruskal-Wallis H-test with Bonferroni correction). **b-c)** Selected Level 2 (bold) and Level 3 (non-bold) KEGG Ortholog pathways enriched in HS (HSN + HSL) and normal skin. Val = valine, leu = leucine, ile = isoleucine. Multiple group statistical analysis performed in STAMP (Kruskal-Wallis H-test with Bonferroni correction). * indicates significance compared to normal (p-value < 0.05).

### Metabolic pathways are influenced by different genera in normal and HS skin

We next identified the specific genera associated with KEGG Orthologs involved in key metabolic pathways. To do this, we combined all significant KEGG Orthologs (K number) affiliated with each pathway and determined the top 10 contributing genera to each pathway using PICRUSt (**Figure 6, Table S9**). The genera contributing to each pathway differed between HS and normal skin. For example, C*utibacterium* contributed significantly to *propanoate metabolism* in normal skin (p=0.004), while *Corynebacterium* was the dominant contributor in HS skin (p=0.062) (**Figure 6a, Table S10**). *Cutibacterium* was the dominant contributor to *retinol metabolism* in normal skin, while no single genera dominated in HS skin (p=0.0003) (**Figure 6b, Table S10**). *Corynebacterium* is a major contributor to multiple amino acid metabolism pathways in HS skin including *valine, leucine, and isoleucine biosynthesis* and *degradation* as well as *arginine and proline metabolism* (**Figure 6c-e, Table S10**). Overall, dysbiosis not only changes which bacteria are present, but also impacts the biological functions of the skin microbial community.

**Figure 6:**
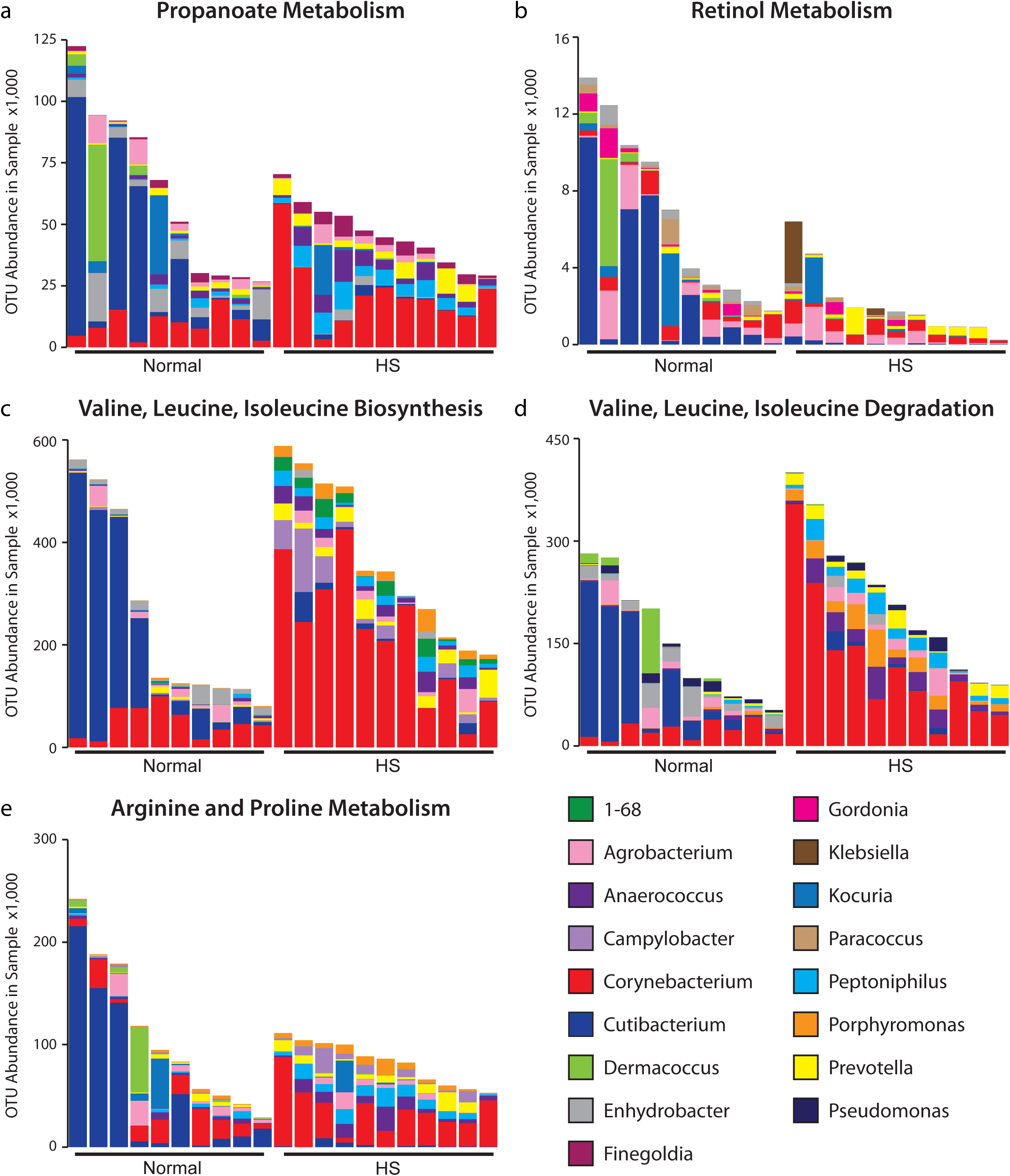
Metabolic pathways are influenced by different genera in normal and HS skin. Top 10 genera contributing to metabolic pathways with each column representing the mean bacterial OTU abundance from all samples collected from one subject. **a)** *Propanoate Metabolism*, **b)** *Retinol Metabolism*, **c)** *Valine, Leucine, Isoleucine Biosynthesis*, **d)** *Valine, Leucine, and Isoleucine Degradation*, **e)** *Arginine and Proline Metabolism*.

## DISCUSSION

Dysbiosis of the skin microbiota is common in many inflammatory skin diseases. Our data are in accordance with previous studies documenting dysbiosis within HS skin compared to normal skin (Guet-Revillet et al. 2017; Nikolakis et al. 2017; Ring et al. 2017b; Ring et al. 2017a). Unlike normal subjects that demonstrate heterogeneity of bacterial communities across body sites, within the follicle and at the skin surface; we demonstrate that this heterogeneity is lost in HS skin. Moreover, we identified metagenomic functional differences between HS and normal skin. Dysbiosis in HS skin may be the result of the disease process, as suggested by increased metabolic turnover, or may contribute to the disease process as suggested by altered amino acid metabolism pathways.

Smoking impacts HS severity and cessation is often recommended to ease symptoms (Sartorius et al. 2009). While differences in β-diversity based on lifestyle choices did not reach statistical significance in our study, smoking and alcohol likely impact the skin microbiome in HS patients. Both smoking and alcohol use increase inflammation (Rom et al. 2013; Szabo and Saha 2015), which may alter skin bacteria composition through host-microbiome crosstalk. While smoking and alcohol use impact β-diversity of the gut microbiome (Capurso and Lahner 2017), to our knowledge ours is the first study to link smoking and alcohol use to an altered skin microbiome.

We uncovered changes in specific bacterial compositions comparing HS and normal skin. *Cutibacterium*, a commensal bacterium found within the PSU, was significantly decreased in HSL and HSN compared with normal skin, consistent with other published work (Ring et al. 2017b). Decreased abundance of *Cutibacterium* in HS is likely due to destruction of the PSU during disease progression (Kamp et al. 2011). Loss of *Cutibacterium* in non-lesional skin of HS subjects suggests that disruption of PSU architecture may precede the development of clinically detectable HS lesions. Alternatively, antimicrobial peptides and inflammatory mediators initiated by follicular keratinocytes in the early stages of HS may form a microenvironment inhospitable to *Cutibacterium* colonization (Hotz et al. 2016). Short-chain fatty acids, including propionic acid, lactic acid, and acetic acid, produced by *Cutibacterium* have antimicrobial activity against *Staphylococcus aureus* and other opportunistic pathogens (Shu et al. 2013). Loss of *Cutibacterium* together with a decrease in *propanoate metabolism* may relate to the observed increased abundance of opportunistic pathogens including *Peptoniphilus, Porphyromonas, Agrobacterium, Pseudomonas*, and *Arcanobacterium* in HS skin. *Peptoniphilus* and *Porphyromonas* have been previously identified in HS lesions (Ring et al. 2017b), and all these pathogens have been isolated from healthy and immunocompromised patients with skin infections, chronic wounds, abscesses, and pressure ulcers (Gellatly and Hancock 2013; Hulse et al. 1993; Kim et al. 2016; Murphy and Frick 2013; Tan et al. 2006). Methods that stabilize the *Cutibacterium* community to maintain homeostasis and keep pathogenic communities at bay may help improve symptoms of HS.

Using metagenomics, we identified several metabolic pathways that are different between HS and normal skin and these differences may impact HS pathology. Amino acid metabolism may impact body odor and skin pH. Malodor significantly impairs the quality of life in HS patients (Alavi et al. 2018). The breakdown of isoleucine to isovaleric acid contributes to “sour” odor (Lam et al. 2018), and we identified altered isoleucine biosynthesis and degradation pathways in HS skin. *Corynebacterium* significantly contributed to these pathways. In other studies, *Corynebacterium* produced thioalcohols (3M3SH, 3M2H, HMHA) which are linked with “sour” malodor in adults (Bawdon et al. 2015; Fredrich et al. 2013; Taylor et al. 2003). Interestingly, in individuals who do not use underarm hygiene products, as some HS patients do not, *Corynebacterium* was the most abundant genera (Urban et al. 2016). Even though *Corynebacterium* is a common skin commensal, in the context of HS, it may trigger insidious inflammatory and odor changes. Altered arginine and proline metabolism in HS skin may incite an acidic skin pH, which may further contribute to the state of dysbiosis.

Finally, our data suggest that rather than targeting the bacterial dysbiosis of HS with antibiotics, there is an opportunity to target the metabolic profiles of bacteria, with the goal of restoring a “normal metabolome.” For example, we observed a significant decrease in retinol metabolism among bacteria in HS skin. Apart from pharmacologic intervention with systemic retinoids, alternative approaches to restore a normal metabolome using prebiotics, probiotics or microbiome transplants may be of interest in HS, similar to the use of skin microbiome transplants in atopic dermatitis (Blok et al. 2013; Nakatsuji et al. 2017). In acne individuals, vitamin B12 supplementation impacted *C. acnes* B12 metabolic pathways suggesting that host interventions can positively or negatively impact the bacterial metabolome (Kang et al. 2015). Influencing the bacterial metabolome through vitamin supplementation in HS skin may be possible as results from this study indicated that several amino acid and vitamin metabolism pathways are altered in HS skin. Computational analyses provide initial clues to understanding the functions of individual bacteria within the microbial community; however future studies confirming these hypotheses are warranted.

In conclusion, bacterial dysbiosis is present in HS compared to normal skin. This dysbiosis is apparent at the skin’s surface, in the follicle, and at distinct body sites. Since dysbiosis can impact the biological function of the skin microbiota, understanding the bacterial metabolome and how it influences the host may be critical to determining the contribution of bacteria to HS pathology and may suggest novel therapeutic options.

## MATERIALS AND METHODS

*Detailed methods are provided in Supplementary Materials.*

### Study participants

Twenty one volunteers (10 normal and 11 HS) were recruited under an approved IRB protocol (Penn State College of Medicine) and provided written informed consent prior to the study. Subjects included males and females, ages 18-65. Subjects were excluded if they had known allergies to cyanoacrylates or had used topical antimicrobials in the previous 2 weeks or systemic antibiotics in the previous month.

### Microbiome sampling and DNA extraction

Two methods were used to sample the skin microbiome of the axilla and groin—cyanoacrylate glue to sample the pore/follicle and swabs to sample the skin surface. Non-lesional and lesional skin were sampled in HS patients, while healthy skin was sampled in normal subjects. Mock community control (MCC; ATCC) (**Figure S3**) and negative sample controls (i.e. no skin contact) were processed alongside samples from human subjects.

### 16S rRNA gene sequencing (V3-V4)

16S rRNA gene sequencing was performed at the Genome Sciences and Bioinformatics Core the Penn State College of Medicine. Sequencing was performed using Illumina MiSeq (paired 300 bp reads) for the V3-V4 region using a two-step, tailed PCR approach that has been validated by Illumina (Illumina, Inc, San Diego, CA).

### Sequence processing and taxonomic mapping

Fastq samples were processed using Pear (version 0.9.6) to join paired end reads, and subsampling was performed at a depth of 75,000 reads using seqtk (version 1.0-r82). Samples that did not meet minimum quality scores or subsampling requirements were removed from analysis (**Table S2**). Operational Taxonomic Units (OTUs) were determined using QIIME (version 1.9.1_py2.7.11) (Caporaso et al. 2010). Reads were mapped with GreenGenes 16S rRNA gene reference database (version 13.8) at 97% sequence similarity (McDonald et al. 2012) using the default OTU picking method, uclust (Edgar 2010). Data was converted to biom format (genus level) and prepared for downstream statistical analyses in R. Genera that had ≥ 2% relative abundance in negative controls were removed from all downstream analyses.

### Diversity analyses and statistics

All data analyses were conducted using the statistical software R. Diversity between samples were compared by calculating both α-diversity (Shannon diversity index – within sample diversity) and β-diversity (Bray-Curtis dissimilarity index – between-sample diversity; 0 = identical communities, 1 = dissimilar communities). Differences in Shannon index between groups were evaluated by the Wilcoxon rank-sum test. Differences in Bray-Curtis dissimilarity index were assessed by MiRKAT (Zhao et al. 2015) or GHT (Zhao et al. 2018) for HSN-HSL paired samples. Both MiRKAT and GHT evaluate the overall shift in microbiome composition. Differential abundances of individual taxon were tested using the Wilcoxon rank sum test followed by the Bonferroni correction to preserve the family-wise error rate (FWER) of 0.05.

### Functional analyses

Functional differences of OTUs were determined using PICRUSt (Langille et al. 2013). Functional data were generated to level 3 functional categories of the Kyoto Encyclopedia of Genes and Genomes (KEGG) orthologs (Kanehisa and Goto 2000). Statistical Analysis of Metagenomic Profile (STAMP) package (Parks et al. 2014) was used to make comparisons between multiple groups (normal, HSN, and HSL) using the Kruskal-Wallis H-test with Bonferonni correction.

## AUTHOR CONTRIBUTIONS

**LEAD AUTHORS**

**\*AMS:** conceptualization, data curation, formal analyses, investigation, software, visualization, and writing.

**\*LCC:** conceptualization, data curation, funding acquisition, methodology, resources, and writing.

**AMN:** conceptualization, data curation, formal analyses, funding acquisition, methodology, project administration, resources, supervision, visualization, and writing.

*equal contributions

## SUPPORTING AUTHORS

**XZ:** formal analyses, software, visualization, and writing.

**KB:** formal analyses, software, and visualization.

**ZC:** investigation and methodology.

**YIK:** methods, investigation, resources, and writing.

**SLG:** project administration, data curation, and resources.

**ALL:** project administration, data curation, and resources.

**JSK:** resources, formal analyses, and writing.

All authors edited the manuscript, gave final approval for publication, and agreed to be accountable for the work.

## ABBREVIATIONS

FWER: family-wise error rate
HS: Hidradenitis Suppurativa
HSN: HS non-lesional
HSL: HS lesional
KEGG: Kyoto Encyclopedia of Genes and Genomes
MCC: mock-community control
MiRKAT: Microbiome Regression Kernel Association Test
OTUs: operational taxonomic units
PICRUSt: Phylogenetic Investigation of Communities by Reconstruction of Unobserved States
PSU: pilosebaceous unit
STAMP: Statistical Analysis of Metagenomic Profile

## ACKNOWLEDGEMENTS

This work was supported by a research grant from the American Acne and Rosacea Society to LCC and the Department of Dermatology Research Endowment at the Penn State College of Medicine. The authors thank Dr. Dave Adams (PSU Dermatology) for assistance with patient recruitment and clinical expertise and Jacob B. Hall, PhD for advice and guidance with data analyses. We appreciate language editing services and writing instruction provided by DerMEDit (www.dermedit.com).

