## Supplemental Files for "Loss of microbial diversity and body site heterogeneity in individuals with Hidradenitis Suppurativa"

#### Supplemental Tables, Figures, and Methods

##### Supplemental Table S1. Subject Demographics.

| Normal Volunteers | Sex (M/F) | Age (yrs) | Ethnicity (Hispanic/Latino vs Non-Hispanic/Latino) | Race | Alcohol Use (Y/N; frequency) | Smoking (Y/N; frequency) | Soap use (Y/N, Brand) | Deodorant (Y/N; Brand) | Systemic Medications (Y/N;drug) | HS Personal Hx (duration yrs; body sites) | HS Hurley Stage | Family Hx of HS (Y/N) | Other Inflammatory Skin conditions (Y/N) |
|---|---|---|---|---|---|---|---|---|---|---|---|---|---|
| 1 | F | 39 | Non-Hispanic | Asian | N | N | Y; Dove | Y; Dove | N | NA | NA | N | N |
| 2 | M | 25 | Non-Hispanic | Asian | Y; casual | N | Y; Bath & Body | Y; Old Spice | N | NA | NA | N | Hx of severe acne |
| 3 | M | 27 | Non-Hispanic | Caucasian | N | N | Y; Up &Up | Y; Old Spice | N | NA | NA | N | N |
| 4 | F | 29 | Non-Hispanic | Caucasian |  | N | Y; Lush | Y; Ban | Y; Symbicort, Nexium | NA | NA | N | Hx of pilonidal cyst, not active |
| 5 | M | 61 | Non-Hispanic | Caucasian | N | N | Y; NA | Y; NA | Y; Lipitor, aspirin | NA | NA | N | N |
| 6 | F | 58 | Non-Hispanic | Caucasian | Y; weekly | N | Y; Goat's milk soap | Y; Dove | Y; Effexor; Celebrex, Albuterol, Flonase; Zyrtec; | NA | NA | N | N |
| 7 | F | 47 | Non-Hispanic | Caucasian | N | N | Y; Ivory, Bath & Body | Y; Sauve | N | NA | NA | N | N |
| 8 | F | 44 | Non-Hispanic | Caucasian | N | N | Y; Bath & Body | Y; Sauve | N | NA | NA | N | N |
| 9 | F | 21 | Non-Hispanic | Caucasian | Y; casual | N | Y; Bath & Body | Y; Dove | Y: tri-sprintec | NA | NA | N | N |
| 10 | M | 19 | Non-Hispanic | Caucasian | N | N | Y; Irish Spring | Y; Old Spice | N | NA | NA | N | N |
| **HS Patients** |  |  |  |  |  |  |  |  |  |  |  |  |  |
| 1 | F | 34 | Non-Hispanic | Caucasian | Y; casual | N | Y; Johnson & Johnson Baby | N | Y; metformin, spironolactone, sovereign silver, emuaid max | 7; axilla, breast, buttock, groin | 2 | Y; Father, Paternal Grandmother | Hx of pilonidal cyst and severe acne, not currently active. |
| 2 | F | 34 | Non-Hispanic | Caucasian | N | N | Y; Method | Y; Tom's of Maine | Y; Flonase, Ibuprofen, Letrozole; past use | 8; axilla, breast, buttock, groin | 3 | N | N |
| 3 | F | 50 | Non-Hispanic | Caucasian | N | N | Y, Emuaid | N | Y; orencia, synthroid, gabapentin, zegerid, diovan, topiramate, nortriptyline, regalin, methotrexate, astepro, fluticasone, folic acid, flexeril, ambien, brintellix, propranolol hcl er, tramadol | 5; axilla, breast, groin | 3 | Y, Mother | N |
| 4 | F | 34 | Non-Hispanic | Black/African American | Y; casual | N | Y, Method | Y; Sauve | Y; Humira | 4; axilla, breast, groin | 1 | N | N |
| 5 | F | 33 | Hispanic | Puerto Rican | Y; casual | Y; 1/3 pack per day (15 yrs) | Y; Dial | N | Y; abilify, cymbalta, ambien, valium | 13; axilla, breast, buttock, groin | 3 | N | N |
| 6 | F | 33 | Non-Hispanic | Caucasian | N | Y; 1-2 packs/day | Y; Dial | N | Y; Welbutrin | 6; axilla, breast, groin | 2 | N | N |
| 7 | F | 36 | Non-Hispanic | Caucasian | Y; daily | N | Y; Hibiclens | N | Y; Humira, Lisinopril | 8; axilla, breast, groin | 3 | N | Hx of severe acne; not active |
| 8 | F | 47 | Non-Hispanic | Caucasian | Y; casual | Y; 8 cigarettes/day (33 yrs) | Y; Shower gel | Y; Dove | Y; losartan /HCT, levothyroxine | 3; axilla, breast, groin | 2 | N | Hx of severe acne; not active |
| 9 | F | 20 | Hispanic | Black/African American | Y; casual | N | Y; Dove | Y; Dove | N | 8; axilla, breast, buttock, groin | 3 | N | N |
| 10 | M | 55 | Non-Hispanic | Caucasian | N | Y; 1 pack/day (29 yrs) | Y; Ivory | N | Y; isosorbide, hydrochlorothiazide, amlodipine, eliquis, aspirin | 4; axilla, buttock, groin | 3 | Y; Father | Hx of severe acne; not active |
| 11 | F | 35 | Non-Hispanic | Caucasian | N | Y; 4 cigarettes/day (22 yrs), Vape Pen (1 yr) | Y; Dove | Y; Dove | N | 1; axilla, breast, buttock, groin | 2 | N | N |

#### Supplemental Table S2. Quality-trimmed and filtered sequence counts for each sample

Forward and Reverse sequences for each sample were joined using Pear and filtered at PHRED score 33. The V3-V4 data set contained 31,611,102 sequence reads with a median of 246,962 sequence reads per sample. **Bolded** samples were not included in analyses after subsampling to 75,000 sequences.

| Normal Sample ID | Sequence Count | Normal Sample ID | Sequence Count | HS Sample ID | Sequence Count | HS Sample ID | Sequence Count |
|---|---|---|---|---|---|---|---|
| N001_Axilla_Glue | 296866 | **N010_Axilla_Glue** | **73642** | HS004_Lesion_Groin_Swab | 226799 | HS009_Non-lesion_Groin_Swab | 664410 |
| N001_Axilla_Swab | 312072 | N010_Axilla_Swab | 118553 | HS005_Non-lesion_Axilla_Glue | 164747 | **HS009_Lesion_Axilla_Glue** | **65428** |
| N001_Groin_Glue | 175148 | **N010_Groin_Glue** | **16866** | HS005_Non-lesion_Axilla_Swab | 342722 | HS009_Lesion_Axilla_Swab | 248066 |
| N001_Groin_Swab | 165463 | N010_Groin_Swab | 101310 | HS005_Non-lesion_Groin_Glue | 214836 | **HS009_Lesion_Groin_Glue** | **15383** |
| **N002_Axilla_Glue** | **43930** | **HS Sample ID** | | HS005_Non-lesion_Groin_Swab | 218240 | HS009_Lesion_Groin_Swab | 1872610 |
| N002_Axilla_Swab | 268401 | HS001_Non-lesion_Axilla_Glue | 1104551 | HS005_Lesion_Axilla_Glue | 178103 | HS010_Non-lesion_Axilla_Glue | 276697 |
| N002_Groin_Glue | 173291 | HS001_Non-lesion_Axilla_Swab | 289970 | HS005_Lesion_Axilla_Swab | 86042 | **HS010_Non-lesion_Axilla_Swab** | **69090** |
| N002_Groin_Swab | 153314 | HS001_Non-lesion_Groin_Glue | 190224 | HS005_Lesion_Groin_Glue | 241333 | HS010_Non-lesion_Groin_Glue | 172314 |
| N003_Axilla_Glue | 293581 | **HS001_Non-lesion_Groin_Swab** | **1115** | HS005_Lesion_Groin_Swab | 258561 | HS010_Non-lesion_Groin_Swab | 132966 |
| N003_Axilla_Swab | 151766 | HS001_Lesion_Axilla_Glue | 618268 | HS006_Non-lesion_Axilla_Glue | 294435 | HS010_Lesion_Axilla_Glue | 150067 |
| N003_Groin_Glue | 247930 | HS001_Lesion_Axilla_Swab | 89895 | HS006_Non-lesion_Axilla_Swab | 119416 | HS010_Lesion_Axilla_Swab | 468254 |
| N003_Groin_Swab | 207352 | HS001_Lesion_Groin_Glue | 272498 | **HS006_Non-lesion_Groin_Glue** | **1042** | HS010_Lesion_Groin_Glue | 282585 |
| N004_Axilla_Glue | 206582 | HS001_Lesion_Groin_Swab | 665789 | HS006_Non-lesion_Groin_Swab | 319596 | HS010_Lesion_Groin_Swab | 98328 |
| N004_Axilla_Swab | 278113 | HS002_Non-lesion_Axilla_Glue | 96425 | **HS006_Lesion_Groin_Glue** | **44966** | HS011_Non-lesion_Axilla_Glue | 87886 |
| **N004_Groin_Glue** | **4058** | HS002_Non-lesion_Axilla_Swab | 259369 | HS006_Lesion_Groin_Swab | 641287 | **HS011_Non-lesion_Axilla_Swab** | **70924** |
| N004_Groin_Swab | 197477 | HS002_Non-lesion_Groin_Glue | 116800 | HS006_Lesion_Axilla_Glue | 316064 | HS011_Non-lesion_Groin_Glue | 146628 |
| N005_Axilla_Glue | 542423 | HS002_Non-lesion_Groin_Swab | 250371 | HS006_Lesion_Axilla_Swab | 795793 | HS011_Non-lesion_Groin_Swab | 198052 |
| N005_Axilla_Swab | 276685 | **HS002_Lesion_Axilla_Glue** | **3987** | HS007_Non-lesion_Axilla_Glue | 253436 | HS011_Lesion_Axilla_Glue | 163797 |
| N005_Groin_Glue | 395472 | HS002_Lesion_Axilla_Swab | 693191 | HS007_Non-lesion_Axilla_Swab | 297845 | HS011_Lesion_Axilla_Swab | 656788 |
| N005_Groin_Swab | 153512 | HS002_Lesion_Groin_Glue | 118027 | HS007_Non-lesion_Groin_Glue | 472296 | HS011_Lesion_Groin_Glue | 135694 |
| N006_Axilla_Glue | 266236 | HS002_Lesion_Groin_Swab | 164175 | HS007_Non-lesion_Groin_Swab | 193866 | HS011_Lesion_Groin_Swab | 225600 |
| N006_Axilla_Swab | 125774 | HS003_Non-lesion_Axilla_Glue | 163401 | HS007_Lesion_Axilla_Glue | 264976 | **Control Samples** | |
| N006_Groin_Glue | 126558 | HS003_Non-lesion_Axilla_Swab | 652023 | HS007_Lesion_Axilla_Swab | 156416 | MCC | 1246037 |
| N006_Groin_Swab | 183011 | HS003_Non-lesion_Groin_Glue | 452064 | HS007_Lesion_Groin_Glue | 342724 | Negative Control 1 | 164523 |
| **N007_Axilla_Glue** | **20067** | HS003_Non-lesion_Groin_Swab | 163106 | HS007_Lesion_Groin_Swab | 488702 | Negative Control 2 | 180046 |
| N007_Axilla_Swab | 155555 | HS003_Lesion_Axilla_Glue | 108189 | HS008_Non-lesion_Axilla_Glue | 203999 | Negative Control 3 | 226978 |
| N007_Groin_Glue | 143535 | HS003_Lesion_Axilla_Swab | 209867 | HS008_Non-lesion_Axilla_Swab | 325385 | Negative Control 4 | 228271 |
| N007_Groin_Swab | 275556 | HS003_Lesion_Groin_Glue | 269557 | HS008_Non-lesion_Groin_Glue | 314396 | | |
| N008_Axilla_Glue | 98487 | HS003_Lesion_Groin_Swab | 148804 | HS008_Non-lesion_Groin_Swab | 261744 | | |
| N008_Axilla_Swab | 172634 | HS004_Non-lesion_Axilla_Glue | 186884 | HS008_Lesion_Axilla_Glue | 75213 | | |
| **N008_Groin_Glue** | **32495** | HS004_Non-lesion_Axilla_Swab | 143212 | HS008_Lesion_Axilla_Swab | 390895 | | |
| N008_Groin_Swab | 123135 | HS004_Non-lesion_Groin_Glue | 221019 | HS008_Lesion_Groin_Glue | 113034 | | |
| N009_Axilla_Glue | 134815 | **HS004_Non-lesion_Groin_Swab** | **994** | HS008_Lesion_Groin_Swab | 245015 | | |
| N009_Axilla_Swab | 160545 | HS004_Lesion_Axilla_Glue | 139042 | **HS009_Non-lesion_Axilla_Glue** | **32541** | | |
| N009_Groin_Glue | 111712 | HS004_Lesion_Axilla_Swab | 312423 | HS009_Non-lesion_Axilla_Swab | 731319 | | |
| N009_Groin_Swab | 147224 | HS004_Lesion_Groin_Glue | 353706 | HS009_Non-lesion_Groin_Glue | 191659 | | |

#### Supplemental Table S3: Distribution of 117 samples for analyses

| Classification | Normal | HS (N + L) | MCC | Negative Controls |
|---|---|---|---|---|
| Axilla, Glue | 7 | 19 (10+9) | --- | --- |
| Axilla, Swab | 10 | 20 (9+11) | --- | --- |
| Groin, Glue | 7 | 19 (10+9) | --- | --- |
| Groin, Swab | 10 | 20 (9+11) | --- | --- |
| Total | 34 | 78 (38+40) | 1 | 4 |

**<u>Supplemental Table S4: Significantly different genera between (A) Normal and HS Non-lesion (B) Normal and HS Lesion</u>.** List of genera that are significantly different between indicated groups: (A) Normal vs HSN or (B) Normal vs HSL under the family-wise error rate of 0.05. The p-value corresponds to the Bonferroni-adjusted p-value. Genera in **BOLD** are in the Top 20 most abundant genera (**Table S4**).

**A:** Normal vs HSN

| | Mean Rel. Abundance % (SD) | | | |
|---|---|---|---|---|
| **Genus** | **Normal** | **HSN** | **Fold Change** | **P-value (FWER-adj)** |
| **Cutibacterium** | 18.84 (0.284) | 1.322 (0.027) | 14.3 | 4.6E-05 |
| **1-68** | 0.123 (0.004) | 1.853 (0.027) | 0.07 | 1.4E-03 |
| **Peptoniphilus** | 1.794 (0.029) | 7.845 (0.068) | 0.23 | 2.2E-03 |
| **Gordonia** | 0.404 (0.008) | 0.038 (0.002) | 10.6 | 2.4E-03 |
| Sphingobacterium | 0.050 (0.001) | 0.001 (0.001) | 50.0 | 4.2E-03 |
| **WAL_1855D** | 0.079 (0.002) | 1.919 (0.027) | 0.04 | 9.0E-03 |
| Leptotrichia | 0.170 (0.004) | 0.004 (0.001) | 42.5 | 1.3E-02 |
| Janthinobacterium | 0.065 (0.002) | 0.002 (0.001) | 32.5 | 1.5E-02 |
| **Porphyromonas** | 0.611 (0.014) | 3.995 (0.047) | 0.15 | 1.6E-02 |
| Roseomonas | 0.142 (0.003) | 0.007 (0.001) | 20.3 | 2.0E-02 |
| Microbacterium | 0.069 (0.002) | 0.018 (0.001) | 3.83 | 2.1E-02 |
| **Enhydrobacter** | 2.291 (0.037) | 0.257 (0.010) | 8.91 | 2.7E-02 |
| Haemophilus | 0.155 (0.002) | 0.008 (0.001) | 19.4 | 2.9E-02 |
| **Anaerococcus** | 2.555 (0.031) | 8.299 (0.075) | 0.31 | 3.3E-02 |
| **Dialister** | 0.192 (0.003) | 1.164 (0.013) | 0.16 | 3.9E-02 |

**B:** Normal vs HSL

| | Mean Rel. Abundance % (SD) | | | |
|---|---|---|---|---|
| **Genus** | **Normal** | **HSL** | **Fold Change** | **P-value (FWER-adj)** |
| **Cutibacterium** | 18.84 (0.281) | 0.711 (0.017) | 26.5 | 9.7E-08 |
| **Peptoniphilus** | 1.794 (0.029) | 10.32 (0.087) | 0.17 | 6.6E-05 |
| **Rothia** | 0.405 (0.011) | 0.013 (0.001) | 31.2 | 6.5E-04 |
| **Porphyromonas** | 0.611 (0.014) | 4.390 (0.051) | 0.14 | 8.5E-04 |
| **Anaerococcus** | 2.555 (0.031) | 9.456 (0.082) | 0.27 | 1.2E-03 |
| **Enhydrobacter** | 2.291 (0.037) | 0.232 (0.011) | 9.88 | 1.5E-03 |
| **WAL_1855D** | 0.079 (0.002) | 1.532 (0.025) | 0.05 | 3.0E-03 |
| Atopobium | 0.014 (0.001) | 0.346 (0.006) | 0.04 | 3.6E-03 |
| **Acinetobacter** | 1.296 (0.016) | 0.426 (0.011) | 3.04 | 5.1E-03 |
| **Neisseria** | 0.429 (0.010) | 0.015 (0.001) | 28.6 | 5.6E-03 |
| Leptotrichia | 0.170 (0.004) | 0.007 (0.001) | 24.3 | 5.8E-03 |
| **Gordonia** | 0.404 (0.008) | 0.011 (0.001) | 36.7 | 6.0E-03 |
| Roseomonas | 0.142 (0.003) | 0.008 (0.001) | 17.8 | 7.7E-03 |
| **1-68** | 0.123 (0.004) | 1.718 (0.025) | 0.07 | 8.1E-03 |
| **Finegoldia** | 2.177 (0.031) | 6.579 (0.064) | 0.33 | 1.1E-02 |
| **Dialister** | 0.192 (0.003) | 1.908 (0.026) | 0.10 | 1.6E-02 |
| Williamsia | 0.141 (0.003) | 0.001 (0.001) | 141 | 1.8E-02 |
| Peptococcus | 0.007 (0.001) | 0.228 (0.004) | 0.03 | 2.0E-02 |
| Delftia | 0.338 (0.009) | 0.061 (0.003) | 5.54 | 2.7E-02 |
| ph2 | 0.035 (0.002) | 0.129 (0.002) | 0.27 | 2.7E-02 |
| **Pseudomonas** | 1.120 (0.018) | 0.298 (0.006) | 3.76 | 4.0E-02 |

**Supplemental Table S5: Relative abundance of the top 20 genera for Normal, HS Non-lesion, and HS Lesion skin.**

| Normal Skin (n=34) | | | HS Non-Lesion (n=36) | | | HS Lesion (n=36) | | |
|---|---|---|---|---|---|---|---|---|
| Genus | Abundance (%) | # of Samples* | Genus | Abundance (%) | # of Samples* | Genus | Abundance (%) | # of Samples* |
| Staphylococcus | 26.2 | 34 | Corynebacterium | 17.3 | 36 | Corynebacterium | 19.4 | 36 |
| Cutibacterium | 18.8 | 34 | Staphylococcus | 17.3 | 36 | Staphylococcus | 14.7 | 36 |
| Corynebacterium | 8.9 | 34 | Anaerococcus | 8.3 | 36 | Peptoniphilus | 10.3 | 36 |
| Anaerococcus | 2.6 | 30 | Peptoniphilus | 7.8 | 36 | Anaerococcus | 9.5 | 36 |
| Enhydrobacter | 2.3 | 29 | Finegoldia | 5.6 | 36 | Finegoldia | 6.6 | 36 |
| Finegoldia | 2.2 | 34 | Prevotella | 4.0 | 31 | Prevotella | 4.6 | 36 |
| Peptoniphilus | 1.8 | 32 | Porphyromonas | 4.0 | 32 | Porphyromonas | 4.4 | 32 |
| Agrobacterium | 1.8 | 34 | Campylobacter | 2.0 | 28 | Dialister | 1.9 | 29 |
| Dermacoccus | 1.5 | 19 | WAL_1855D | 1.9 | 29 | Campylobacter | 1.8 | 28 |
| Prevotella | 1.4 | 26 | 1-68 | 1.9 | 32 | 1-68 | 1.7 | 29 |
| Acineobacter | 1.3 | 34 | Cutibacterium | 1.3 | 36 | WAL_1855D | 1.5 | 31 |
| Pseudomonas | 1.1 | 33 | Dialister | 1.2 | 29 | Fusobacterium | 1.4 | 25 |
| Lactobacillus | 1.0 | 23 | Fusobacterium | 1.1 | 20 | Parvimonas | 1.0 | 15 |
| Kocuria | 0.9 | 25 | Agrobacterium | 1..0 | 32 | Cutibacterium | 0.7 | 35 |
| Porphyromonas | 0.6 | 23 | Parvimonas | 0.5 | 16 | Peptostreptococcus | 0.6 | 20 |
| Clostridium | 0.5 | 23 | Bifidobacterium | 0.5 | 18 | Kocuria | 0.6 | 15 |
| Neisseria | 0.4 | 21 | Brevibacterium | 0.5 | 26 | Brevibacterium | 0.6 | 20 |
| Rothia | 0.4 | 26 | Acinetobacter | 0.5 | 29 | Agrobacterium | 0.6 | 30 |
| Gordonia | 0.4 | 19 | Pseudomonas | 0.5 | 32 | Gallicola | 0.4 | 22 |
| Unclassified | 17.5 | 34 | Unclassified | 15.6 | 36 | Unclassified | 11.2 | 36 |

*number of samples in which genera was detected.

**Supplemental Table S6: Aggregated Subject demographics for Metavariable analyses.**

| Demographic | Normal Subjects | HS Subjects |
|---|---|---|
| Total analyzed | 10 | 11 |
| Hurley Stage (I/II/III) | NA | 1/4/6 |
| Age, mean yrs. (SD) | 37 (15.12) | 37.36 (9.72) |
| Female/Male | 6/4 | 10/1 |
| Caucasian/not Caucasian | 7/3 | 8/3 |
| Alcohol use | 3 | 6 |
| Smoking use* | 0 | 5 |
| Deodorant use** | 10 | 5 |
| Medication use | 4 | 9 |

*p=0.035 (Fisher Exact Test)
**p=0.023(Fisher Exact Test)

**Supplemental Table S7: Top 20 most abundant genera identified in each Hurley Stage Classification**

| Stage I (n = 1) | | Stage II (n = 4) | | Stage III (n = 6) | |
|---|---|---|---|---|---|
| Genus | Abundance (%) | Genus | Abundance (%) | Genus | Abundance (%) |
| Staphylococcus | 22.8 | Staphylococcus | 23.1 | Corynebacterium | 19.2 |
| Corynebacterium | 12.0 | Corynebacterium | 16.5 | Staphylococcus | 12.2 |
| Prevotella | 9.5 | Anaerococcus | 10.0 | Peptoniphilus | 9.9 |
| Finegoldia | 7.4 | Peptoniphilus | 7.7 | Anaerococcus | 8.4 |
| Peptoniphilus | 6.2 | Finegoldia | 4.1 | Finegoldia | 6.8 |
| Anaerococcus | 5.0 | Porphyromonas | 3.0 | Prevotella[1] | 6.2 |
| Dialister | 2.7 | Campylobacter | 2.3 | Porphyromonas | 5.8 |
| WAL_1855D | 2.5 | 1-68 | 1.6 | Fusobacterium | 2.7 |
| Campylobacter | 2.5 | Propionibacterium | 1.6 | WAL_1855D | 2.1 |
| Propionibacterium | 2.2 | Agrobacterium | 1.3 | 1-68 | 2.1 |
| Porphyromonas | 0.7 | WAL_1855D | 1.2 | Dialister | 1.7 |
| Peptostreptococcus | 0.4 | Dialister | 1.1 | Parvimonas | 1.6 |
| Mobiluncus | 0.4 | Prevotella[1] | 1.0 | Campylobacter | 1.2 |
| Pseuoclavibacter | 0.3 | Kocuria | 0.8 | Peptostreptococcus | 0.6 |
| 1-68 | 0.3 | Bifidobacterium | 0.7 | Gemella | 0.6 |
| Varibaculum | 0.3 | Acinobacter | 0.7 | Atopobium[2] | 0.6 |
| ph2 | 0.2 | Brevibacterium | 0.6 | Agrobacterium | 0.5 |
| Atopobium | 0.1 | Pseudomonas | 0.6 | Gallicola | 0.5 |
| Dermabacter | 0.1 | Clostridium | 0.4 | Brevibacterium | 0.5 |
| Unclassified | 23.8 | Unclassified | 16.0 | Unclassified | 10.0 |

Stage II vs Stage III:
[1]**Prevotella** (stage II average abundance = 1.04%, stage III average abundance = 5.93%, adjusted p-value = 0.0003)
[2]**Atopobium** (stage II average abundance = 0.05%, stage III average abundance = 0.51%, adjusted p-value = 0.0009)

### Supplemental Table S8: Significant KEGG Orthologs

| Level 1 | p-values (corrected) | Level 2 | p-values (corrected) | Level 3 | p-values (corrected) | Normal: mean rel. freq. (%) | Normal: std. dev. (%) | HSN: mean rel. freq. (%) | HSN: std. dev. (%) | HSL: mean rel. freq. (%) | HSL: std. dev. (%) |
|---|---|---|---|---|---|---|---|---|---|---|---|
| Cellular Processes | 4.60757 | **Cell Growth and Death** | **0.00319** | Apoptosis | 0.01323 | 0.01203 | 0.01221 | 0.00509 | 0.00783 | 0.00362 | 0.00665 |
| | | | | Cell cycle - Caulobacter | 0.00015 | 0.52671 | 0.04597 | 0.59945 | 0.07780 | 0.61960 | 0.07643 |
| | | | | p53 signaling pathway | 0.04206 | 0.00872 | 0.01037 | 0.00424 | 0.00701 | 0.00303 | 0.00577 |
| | | **Transport and Catabolism** | **0.00048** | Endocytosis | 0.00016 | 0.00145 | 0.00175 | 0.00027 | 0.00072 | 0.00023 | 0.00067 |
| **Environmental Information Processing** | **0.00815** | **Signal Transduction** | **0.04991** | | | | | | | | |
| **Genetic Information Processing** | **1.83E-07** | **Folding, Sorting and Degradation** | **6.52E-07** | Protein export | 0.00732 | 0.77403 | 0.09206 | 0.84280 | 0.09706 | 0.86348 | 0.06231 |
| | | | | RNA degradation | 0.01833 | 0.55167 | 0.03842 | 0.58232 | 0.05121 | 0.59912 | 0.04485 |
| | | **Replication and Repair** | **3.54E-06** | Chromosome | 0.00038 | 1.42112 | 0.16156 | 1.61166 | 0.14509 | 1.64266 | 0.16677 |
| | | | | DNA repair and recombination proteins | 0.00054 | 3.27042 | 0.25787 | 3.56627 | 0.35981 | 3.71475 | 0.32257 |
| | | | | DNA replication | 1.53E-05 | 0.74320 | 0.08679 | 0.85599 | 0.10920 | 0.89515 | 0.09776 |
| | | | | DNA replication proteins | 4.56E-05 | 1.18192 | 0.12397 | 1.36628 | 0.17542 | 1.42609 | 0.19467 |
| | | | | Homologous recombination | 0.00022 | 1.01675 | 0.10594 | 1.15479 | 0.16049 | 1.21892 | 0.15688 |
| | | | | Mismatch repair | 8.14E-07 | 0.82565 | 0.08624 | 0.99559 | 0.14596 | 1.05090 | 0.15439 |
| | | | | Nucleotide excision repair | 0.01022 | 0.44045 | 0.05688 | 0.50233 | 0.09129 | 0.53276 | 0.08092 |
| | | **Replication, recombination and repair proteins** | **1.37E-07** | | | | | | | | |
| | | Transcription | 1.09593 | RNA polymerase | 0.00033 | 0.20587 | 0.02832 | 0.23103 | 0.03022 | 0.23981 | 0.02295 |
| | | | | Transcription machinery | 0.00330 | 0.78757 | 0.07913 | 0.88894 | 0.12105 | 0.91477 | 0.10939 |
| | | **Translation** | **0.00033** | Aminoacyl-tRNA biosynthesis | 0.00028 | 1.37115 | 0.14030 | 1.58385 | 0.23958 | 1.67055 | 0.22546 |
| | | | | Ribosome | 0.00226 | 2.67644 | 0.32799 | 3.08289 | 0.48161 | 3.25351 | 0.46115 |
| | | | | Ribosome Biogenesis | 0.01226 | 1.48909 | 0.09557 | 1.63759 | 0.18325 | 1.71121 | 0.20620 |
| | | | | Ribosome biogenesis in eukaryotes | 0.00051 | 0.06365 | 0.00917 | 0.07418 | 0.01131 | 0.07735 | 0.00932 |
| | | | | Translation factors | 3.57E-06 | 0.53541 | 0.05257 | 0.65200 | 0.11083 | 0.68949 | 0.11627 |
| | | **Translation proteins** | **2.04E-06** | | | | | | | | |
| **Metabolism** | **2.48E-07** | **Amino Acid Metabolism** | **0.00021** | Amino acid related enzymes | 0.00750 | 1.57605 | 0.11304 | 1.70744 | 0.17333 | 1.76282 | 0.14352 |
| | | | | Arginine and proline metabolism | 0.00011 | 1.27063 | 0.10951 | 1.15318 | 0.10608 | 1.13323 | 0.10169 |
| | | | | Tyrosine metabolism | 0.00580 | 0.51563 | 0.04963 | 0.45813 | 0.06130 | 0.44251 | 0.07400 |
| | | | | Valine, leucine and isoleucine biosynthesis | 0.03707 | 0.90063 | 0.07327 | 0.77879 | 0.14634 | 0.74147 | 0.18519 |
| | | Biosynthesis of Other Secondary Metabolites | 0.39045 | Penicillin and cephalosporin biosynthesis | 3.76E-05 | 0.04608 | 0.01025 | 0.03174 | 0.01602 | 0.02689 | 0.01478 |
| | | **Carbohydrate Metabolism** | **3.35E-05** | C5-Branched dibasic acid metabolism | 0.01882 | 0.39979 | 0.05273 | 0.32370 | 0.09131 | 0.29580 | 0.11160 |
| | | | | Citrate cycle (TCA cycle) | 9.35E-05 | 1.08103 | 0.14365 | 0.90217 | 0.13896 | 0.85915 | 0.15890 |
| | | | | Glyoxylate and dicarboxylate metabolism | 0.00019 | 0.68194 | 0.08991 | 0.58938 | 0.10497 | 0.55517 | 0.09552 |
| | | | | Inositol phosphate metabolism | 0.00427 | 0.24633 | 0.09560 | 0.16714 | 0.05013 | 0.15844 | 0.04220 |
| | | | | Pentose and glucuronate interconversions | 2.69E-05 | 0.45239 | 0.07608 | 0.36909 | 0.10106 | 0.33915 | 0.06152 |
| | | **Energy Metabolism** | **0.02251** | Carbon fixation in photosynthetic organisms | 3.75E-05 | 0.58709 | 0.03586 | 0.67125 | 0.07802 | 0.68717 | 0.08681 |
| | | | | Methane metabolism | 0.04657 | 1.16404 | 0.08479 | 1.32008 | 0.19123 | 1.37320 | 0.22598 |
| | | **Enzyme Families** | **1.11E-06** | Peptidases | 3.16E-06 | 1.74116 | 0.11043 | 2.04876 | 0.27338 | 2.13434 | 0.27781 |
| | | Glycan Biosynthesis and Metabolism | 1.37500 | Glycosphingolipid biosynthesis - lacto and neolacto series | 0.00170 | 0.00031 | 0.00048 | 0.00004 | 0.00012 | 0.00001 | 0.00003 |
| | | | | Peptidoglycan biosynthesis | 0.00218 | 0.91700 | 0.12479 | 1.05158 | 0.13483 | 1.09105 | 0.14169 |
| | | **Lipid Metabolism** | **4.56E-05** | | | | | | | | |
| | | Metabolism of Cofactors and Vitamins | 25.64977 | One carbon pool by folate | 0.00705 | 0.66138 | 0.07527 | 0.74478 | 0.11294 | 0.78103 | 0.10199 |
| | | | | Retinol metabolism | 0.00301 | 0.11918 | 0.03306 | 0.07714 | 0.05115 | 0.06321 | 0.05143 |
| | | | | Riboflavin metabolism | 0.00059 | 0.33399 | 0.02127 | 0.35518 | 0.02638 | 0.36860 | 0.02692 |
| | | | | Thiamine metabolism | 0.01193 | 0.57120 | 0.09155 | 0.65384 | 0.07767 | 0.67452 | 0.08407 |
| | | Metabolism of Other Amino Acids | 55.80846 | D-Glutamine and D-glutamate metabolism | 2.53E-06 | 0.16136 | 0.01657 | 0.19621 | 0.03002 | 0.20889 | 0.03073 |
| | | | | Selenocompound metabolism | 0.03173 | 0.45996 | 0.06565 | 0.51635 | 0.05069 | 0.52451 | 0.04552 |
| | | Metabolism of Terpenoids and Polyketides | 4.52620 | Biosynthesis of siderophore group nonribosomal peptides | 4.38E-05 | 0.07462 | 0.01642 | 0.05840 | 0.02988 | 0.05454 | 0.01012 |
| | | | | Prenyltransferases | 0.00637 | 0.47446 | 0.06025 | 0.51244 | 0.06198 | 0.53150 | 0.04234 |
| | | | | Terpenoid backbone biosynthesis | 0.00450 | 0.70408 | 0.06541 | 0.77345 | 0.09546 | 0.80788 | 0.07744 |
| | | | | Zeatin biosynthesis | 0.00976 | 0.04955 | 0.00703 | 0.06073 | 0.01646 | 0.06529 | 0.01656 |
| | | **Nucleotide Metabolism** | **3.84E-05** | Purine metabolism | 2.38E-05 | 2.71613 | 0.22590 | 3.02326 | 0.31792 | 3.14122 | 0.24903 |
| | | | | Pyrimidine metabolism | 0.00117 | 1.96140 | 0.22352 | 2.25573 | 0.32968 | 2.37679 | 0.32774 |
| | | **Xenobiotics Biodegradation and Metabolism** | **0.00780** | Bisphenol degradation | 0.00053 | 0.13242 | 0.04348 | 0.09038 | 0.03069 | 0.08252 | 0.02675 |
| | | | | Drug metabolism - cytochrome P450 | 0.00247 | 0.17097 | 0.06722 | 0.10457 | 0.07110 | 0.08415 | 0.06774 |
| | | | | Metabolism of xenobiotics by cytochrome P450 | 0.00392 | 0.16276 | 0.06312 | 0.10197 | 0.06888 | 0.08187 | 0.06504 |
| | | | | Naphthalene degradation | 0.00178 | 0.32059 | 0.07835 | 0.23468 | 0.08819 | 0.20810 | 0.09102 |
| | | | | Toluene degradation | 4.26E-08 | 0.24978 | 0.03085 | 0.18905 | 0.03733 | 0.17572 | 0.04604 |

*Pathways identifying Level 1 Human Diseases and Organismal Systems and the sub-levels below were removed from the table. These do not accurately describe the activity of the bacteria and are more likely a result of homology between a human gene and bacterial gene with no indication of bacterial function.

**Supplemental Table S9: significant K numbers associated with metabolism pathways in figure 6**

| | | | | | |
|---|---|---|---|---|---|
| **Propanoate Metabolism** | | | | | |
| K00016 | K00249 | K01034 | K01825 | K01913 | K05606 |
| K00114 | K00625 | K01035 | K01847 | K01961 | K07250 |
| K00128 | K00626 | K01505 | K01848 | K01962 | K11264 |
| K00140 | K00656 | K01574 | K01849 | K01963 | K11942 |
| K00169 | K00822 | K01578 | K01895 | K01965 | K13524 |
| K00170 | K00823 | K01659 | K01902 | K01966 | K13788 |
| K00171 | K00925 | K01692 | K01903 | K02160 | K13923 |
| K00172 | K00932 | K01720 | K01905 | K03416 | |
| K00224 | K01026 | K01782 | K01908 | K03417 | |
| **Retinol Metabolism** | | | | | |
| K00001 | K00121 | K09516 | K11147 | K13953 | |
| **Valine, Leucine, and Isoleucine Biosynthesis** | | | | | |
| K00052 | K00163 | K01649 | K01703 | K01870 | K11258 |
| K00053 | K00263 | K01652 | K01704 | K01873 | K14260 |
| K00161 | K00826 | K01653 | K01754 | K09011 | |
| K00162 | K00835 | K01687 | K01869 | | |
| **Valine, Leucine, and Isoleucine Degradation** | | | | | |
| K00020 | K00188 | K00632 | K01641 | K01965 | K09478 |
| K00128 | K00248 | K00822 | K01692 | K01966 | K09699 |
| K00140 | K00249 | K00826 | K01782 | K01968 | K11381 |
| K00166 | K00253 | K01027 | K01825 | K01969 | K11410 |
| K00167 | K00263 | K01028 | K01847 | K05606 | K11942 |
| K00186 | K00382 | K01029 | K01848 | K07250 | K13524 |
| K00187 | K00626 | K01640 | K01849 | K07508 | K13766 |
| **Arginine and Proline Metabolism** | | | | | |
| K00128 | K00611 | K00930 | K01480 | K01941 | K10795 |
| K00137 | K00613 | K00931 | K01484 | K02626 | K10796 |
| K00145 | K00619 | K00934 | K01485 | K03343 | K11358 |
| K00147 | K00620 | K01259 | K01571 | K05526 | K12251 |
| K00260 | K00657 | K01270 | K01572 | K05597 | K12252 |
| K00261 | K00673 | K01425 | K01573 | K06447 | K12253 |
| K00262 | K00797 | K01426 | K01581 | K08687 | K12254 |
| K00273 | K00802 | K01428 | K01583 | K08688 | K12255 |
| K00274 | K00811 | K01429 | K01584 | K09065 | K12256 |
| K00286 | K00812 | K01430 | K01585 | K09251 | K12658 |
| K00294 | K00813 | K01438 | K01611 | K09470 | K12659 |
| K00296 | K00818 | K01457 | K01625 | K09471 | K13043 |
| K00316 | K00819 | K01470 | K01750 | K09472 | K13746 |
| K00318 | K00821 | K01473 | K01755 | K09473 | K13747 |
| K00472 | K00824 | K01474 | K01777 | K10536 | K13821 |
| K00491 | K00840 | K01476 | K01915 | K10793 | K14048 |
| K00542 | K00926 | K01478 | K01940 | K10794 | |

**Supplemental Table S10: Significantly different genera contributions by metabolic pathways in figure 6.**

**(A)** Retinol Metabolism, **(B)** Propanoate Metabolism, **(C)** Valine, Leucine, Isoleucine Biosynthesis**, (D)** Valine, Leucine, Isoleucine Degradation, and **(E)** Arginine and Proline Metabolism. Genera in **BOLD** are contributing to the specific metabolic pathways at significantly different abundance.

A. Propanoate Metabolism

| | **Mean OTU Abundance (SD)** | | | |
|---|---|---|---|---|
| **Genus** | **Normal** | **HS** | **Fold Change (N/HS)** | **P-value (FWER-adj)** |
| Corynebacterium | 9408.432 (5565.696) | 21918.027 (14348.178) | 0.43 | 6.19E-02 |
| **Cutibacterium** | 27828.986 (35536.528) | 875.9 (1261.287) | 31.77 | 3.80E-03 |
| **Enhydrobacter** | 6912.9 (5733.775) | 912.483 (1561.274) | 7.58 | 1.06E-02 |
| **Peptoniphilus** | 1065.718 (1130.401) | 5579.725 (3103.049) | 0.19 | 1.08E-03 |
| Anaerococcus | 1541.213 (1185.038) | 4955.275 (3587.778) | 0.31 | 6.19E-02 |
| Kocuria | 4250.397 (9891.495) | 2120.21 (5989.399) | 2.00 | 1.00E+00 |
| Dermacoccus | 5748.733 (14718.353) | 133.875 (202.188) | 42.94 | 1.00E+00 |
| Prevotella | 1319.125 (890.071) | 4050.622 (3200.201) | 0.33 | 3.57E-01 |
| Agrobacterium | 3385.193 (4082.348) | 1893.971 (2251.154) | 1.79 | 1.00E+00 |
| **Finegoldia** | 1341.352 (1264.477) | 3659.407 (2170.232) | 0.37 | 4.79E-02 |

B. Retinol Metabolism

| | **Mean OTU Abundance (SD)** | | | |
|---|---|---|---|---|
| **Genus** | **Normal** | **HS** | **Fold Change (N/HS)** | **P-value (FWER-adj)** |
| **Cutibacterium** | 3057.617 (3955.067) | 93.196 (128.784) | 32.81 | 3.80E-03 |
| Agrobacterium | 754.917 (919.237) | 426.775 (510.163) | 1.77 | 1.00E+00 |
| Corynebacterium | 578.573 (439.473) | 483.312 (306.107) | 1.20 | 1.00E+00 |
| Kocuria | 500.106 (1163.781) | 249.475 (704.737) | 2.00 | 1.00E+00 |
| Dermacoccus | 676.283 (1731.451) | 15.75 (23.787) | 42.94 | 1.00E+00 |
| Prevotella | 130.338 (109.784) | 344.266 (386.964) | 0.38 | 6.10E-01 |
| Gordonia | 362.513 (512.502) | 98.909 (205.665) | 3.67 | 4.45E-01 |
| Paracoccus | 315.417 (393.519) | 81.691 (99.290) | 3.86 | 7.20E-01 |
| **Enhydrobacter** | 345.779 (286.799) | 45.666 (78.145) | 7.57 | 1.06E-02 |
| Klebsiella | 2.1 (4.483) | 322.726 (960.186) | 0.01 | 1.00E+00 |

C. Valine, Leucine, and Isoleucine Biosynthesis

| | Mean OTU Abundance (SD) | | | |
|---|---|---|---|---|
| **Genus** | **Normal** | **HS** | **Fold Change (N/HS)** | **P-value (FWER-adj)** |
| **Corynebacterium** | 48782.617 (29752.608) | 218839.53 (128347.860) | 0.22 | 1.10E-02 |
| **Cutibacterium** | 166057.833 (203010.341) | 10848.758 (17070.401) | 15.31 | 2.76E-02 |
| **Campylobacter** | 1712.275 (1965.597) | 29916.727 (36557.682) | 0.06 | 4.79E-02 |
| **Prevotella** | 3315.7 (3671.219) | 22052.5 (16192.082) | 0.15 | 4.79E-02 |
| Agrobacterium | 11895.142 (14057.453) | 11561.076 (13898.761) | 1.03 | 1.00E+00 |
| **Anaerococcus** | 2516.7 (1938.988) | 17866.13 (12560.355) | 0.14 | 1.08E-03 |
| **Peptoniphilus** | 1691.7 (2174.949) | 16514.286 (9524.814) | 0.10 | 2.27E-04 |
| **1-68** | 548.861 (998.927) | 17348.201 (13183.768) | 0.03 | 5.50E-03 |
| **Enhydrobacter** | 15280.4 (11872.647) | 3224.727 (5559.133) | 4.74 | 1.72E-02 |
| **Porphyromonas** | 1260.833 (1400.335) | 15818.667 (12150.286) | 0.08 | 5.00E-03 |

D. Valine, Leucine, and Isoleucine Degradation

| | Mean OTU Abundance (SD) | | | |
|---|---|---|---|---|
| **Genus** | **Normal** | **HS** | **Fold Change (N/HS)** | **P-value (FWER-adj)** |
| **Corynebacterium** | 23322.34 (12361.530) | 123073.455 (97696.275) | 0.19 | 2.55E-03 |
| **Cutibacterium** | 73347.933 (88787.727) | 4910.667 (7948.472) | 14.94 | 2.76E-02 |
| **Anaerococcus** | 2636.257 (1945.648) | 19692.39 (14248.084) | 0.13 | 6.80E-04 |
| **Porphyromonas** | 1559.7 (1759.452) | 19773.333 (15187.857) | 0.08 | 5.50E-03 |
| Agrobacterium | 10613.508 (12316.613) | 10149.008 (12135.573) | 1.05 | 1.00E+00 |
| **Enhydrobacter** | 17184.3 (13353.747) | 3624.273 (6245.909) | 4.74 | 1.72E-02 |
| **Peptoniphilus** | 1366.486 (1657.236) | 15865.636 (9807.426) | 0.09 | 3.97E-04 |
| **Prevotella** | 1774.65 (1366.675) | 12198.455 (8611.954) | 0.15 | 2.76E-02 |
| Pseudomonas | 7083.783 (5667.630) | 5563.917 (6345.149) | 1.27 | 1.00E+00 |
| Dermacoccus | 12485.75 (29355.052) | 292.091 (463.765) | 42.75 | 1.00E+00 |

E. Arginine and Proline Metabolism

| | Mean OTU Abundance (SD) | | | |
|---|---|---|---|---|
| **Genus** | **Normal** | **HS** | **Fold Change (N/HS)** | **P-value (FWER-adj)** |
| **Cutibacterium** | 61152.333 (79101.343) | 1863.914 (2575.671) | 32.81 | 3.80E-03 |
| Corynebacterium | 16647.468 (10368.922) | 35985.856 (21710.680) | 0.46 | 1.59E-01 |
| **Anaerococcus** | 2698.479 (2025.299) | 8900.884 (6577.239) | 0.30 | 4.79E-02 |
| **Peptoniphilus** | 1944.271 (2102.271) | 9469.681 (4960.431) | 0.21 | 1.08E-03 |
| Agrobacterium | 7195.666 (8702.778) | 4037.829 (4809.614) | 1.78 | 1.00E+00 |
| Kocuria | 6500.997 (15128.670) | 3242.922 (9160.916) | 2.00 | 1.00E+00 |
| Prevotella | 2352.125 (2081.141) | 6697.76 (5217.008) | 0.35 | 2.95E-01 |
| Dermacoccus | 7777.833 (19913.490) | 181.125 (273.548) | 42.94 | 1.00E+00 |
| Campylobacter | 835.367 (822.799) | 6421.468 (7043.269) | 0.13 | 2.95E-01 |
| **Porphyromonas** | 1245.478 (1212.956) | 5942.546 (4301.281) | 0.21 | 2.06E-02 |

**Supplemental Figure 1: Positive Mock Community Control**

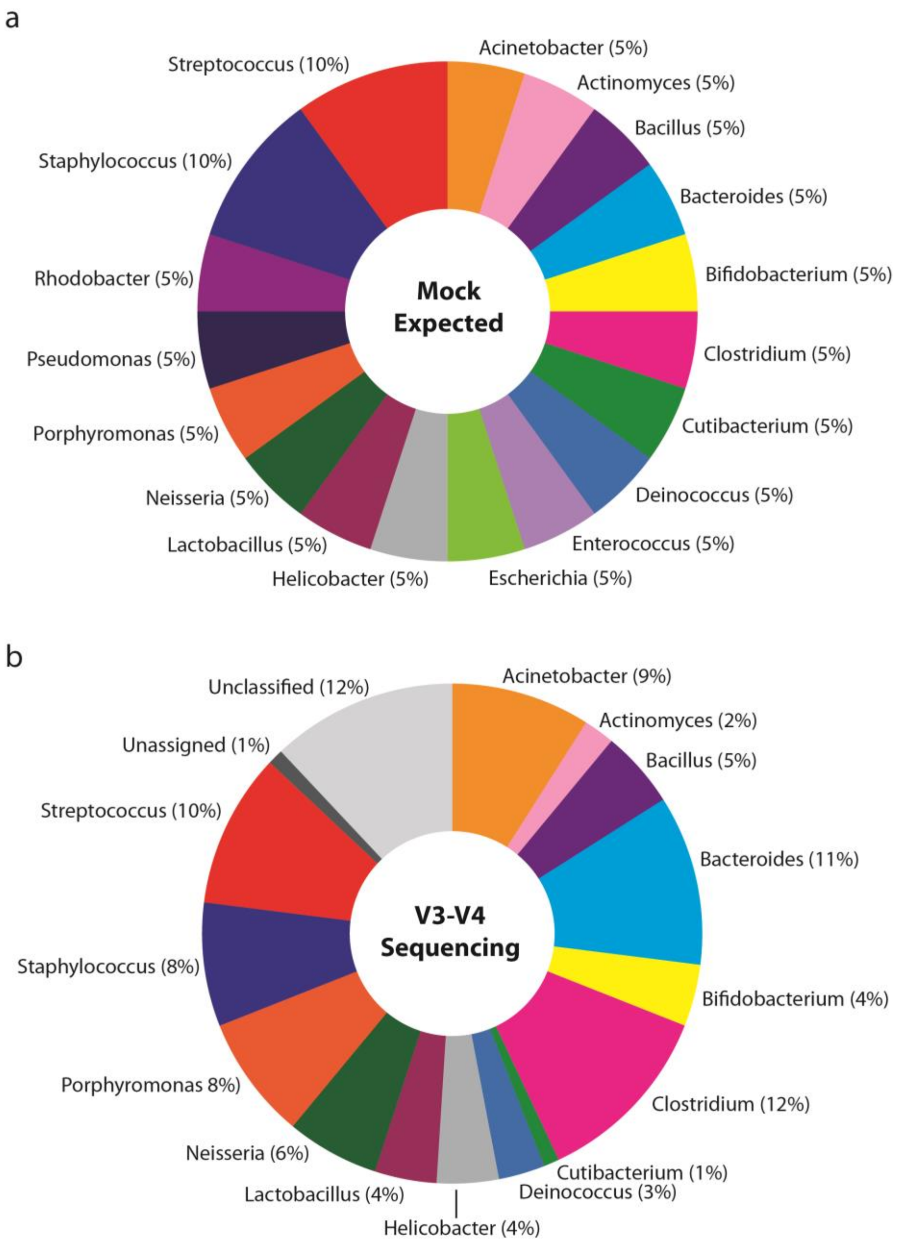

Genus level detection of the positive mock community control **a)** expected percentages **b)** actual sequenced mock community control with 16S rRNA gene V3-V4 sequencing.

__**Supplemental Figure 2: Body site heterogeneity is lost in HS skin compared to normal skin.**__

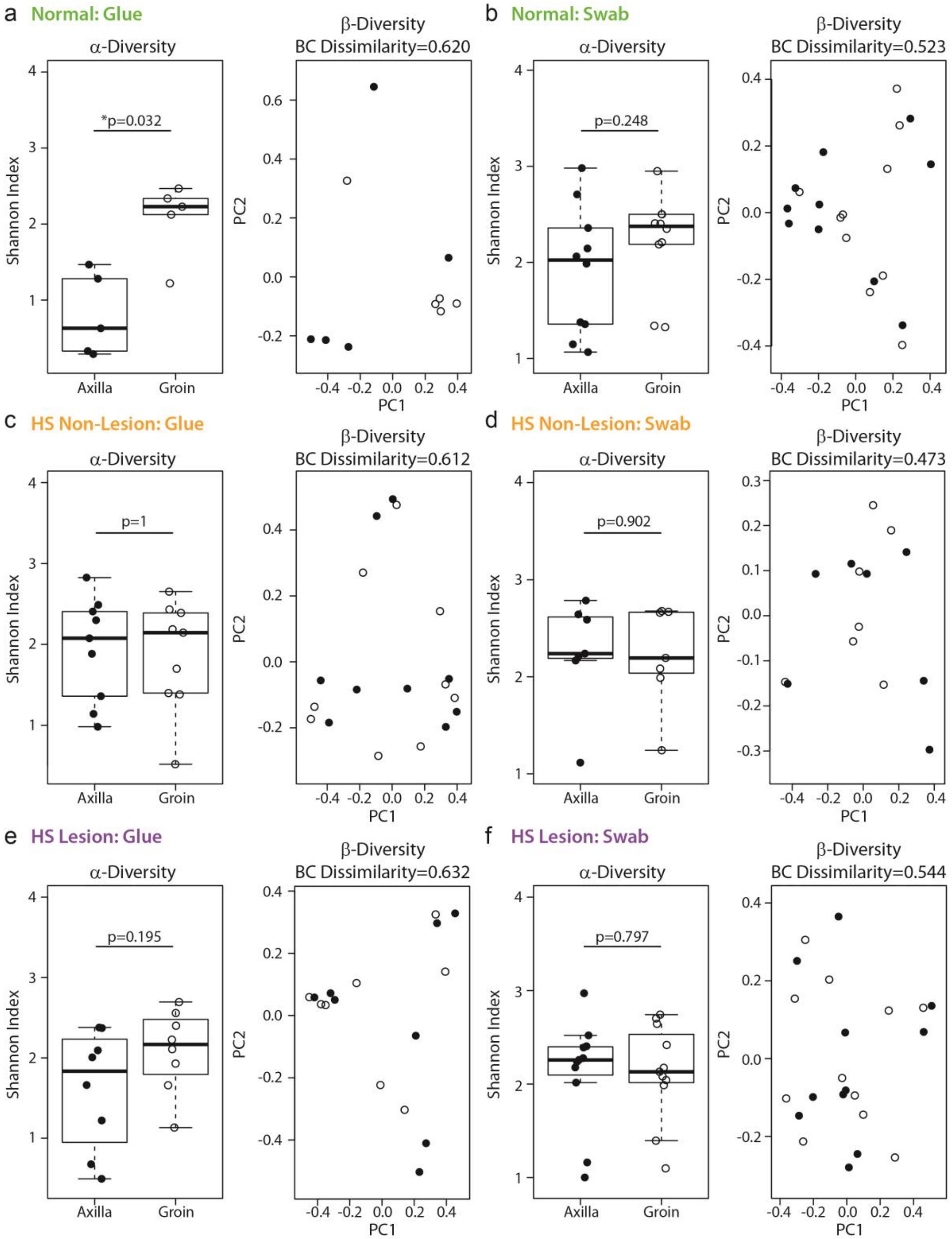

**a-f)** Shannon Diversity Index (α-diversity) and Bray-Curtis Dissimilarity Index (β-diversity) comparison between axilla and groin body sites stratified by skin type and sampling method as indicated. The Wilcoxon Rank Sum test was applied to determine statistical differences in Shannon Index between groups, p values indicated in graph. All analysis were done with paired samples, each sampling method had a matching axilla and groin by disease state **a)** normal glue n=5, **b)** normal swab n=10, **c)** HS non-lesion glue n=9, **d)** HS non-lesion swab n=7, **e)** HS lesion glue n=8, **f)** HS lesion swab n=11.

**<u>Supplemental Figure 3: The follicle and surface microbiome signatures are similar in HS skin.</u>**

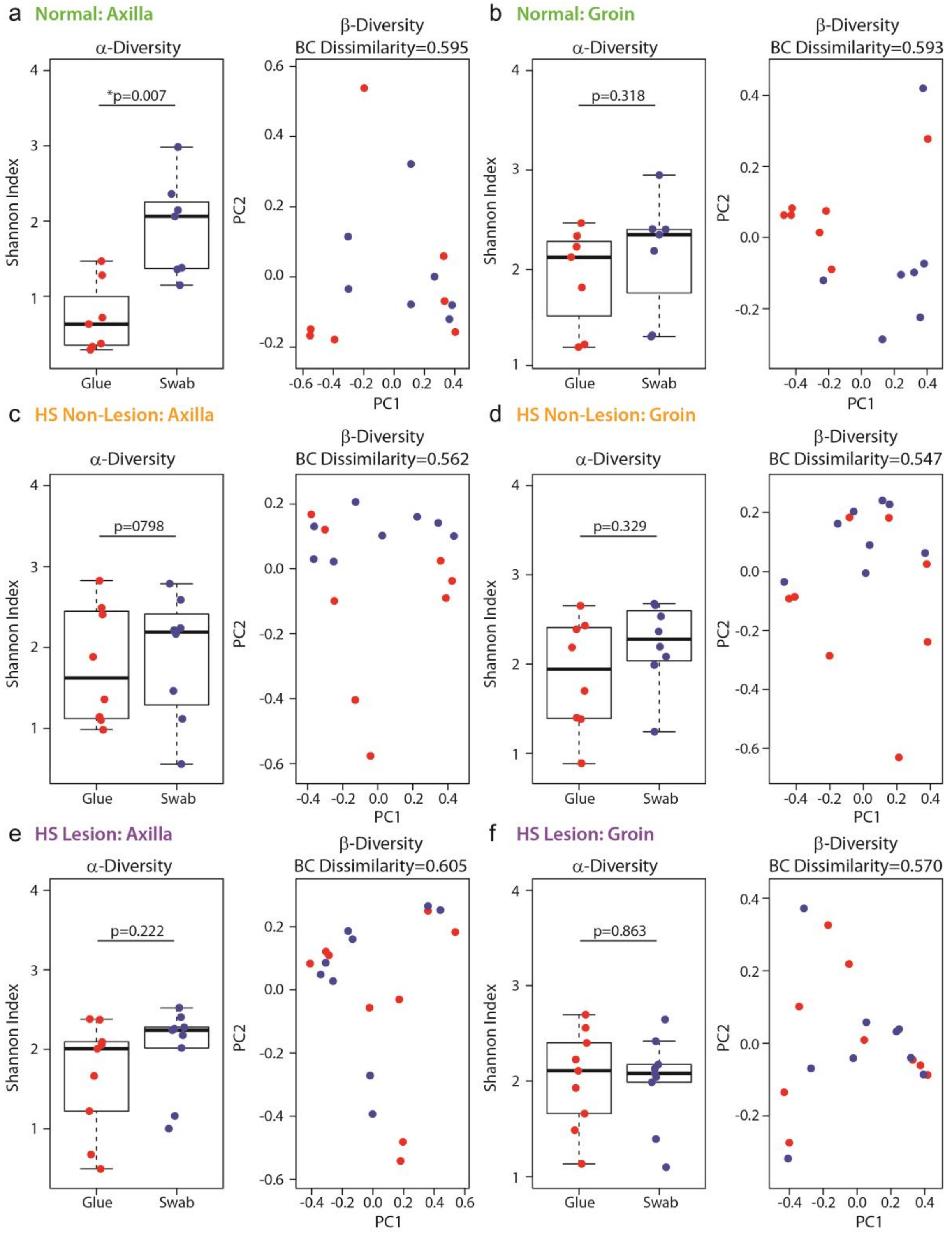

**a-f)** Shannon Diversity Index (α-diversity) and Bray-Curtis Dissimilarity Index (β-diversity) comparison between sampling methods (glue vs swab) stratified by skin type and body site as indicated. The Wilcoxon Rank Sum test was applied to determine statistical differences in Shannon Index between groups, p-values indicated in graph. All analysis were done with paired samples each body site had a matching sampling method by disease state **a)** normal axilla n=7, **b)** normal groin n=7, **c)** HS non-lesion axilla n=8, **d)** HS non-lesion groin n=8, **e)** HS lesion axilla n=9, **f)** HS lesion groin n=9.

#### SUPPLEMENTAL METHODS

##### Supplemental Methods M1. Microbiome sampling methods.

Human Subjects

Ten normal volunteers and 11 HS patients were recruited under an approved Institutional Review Board protocol (#4526; The Pennsylvania State University College of Medicine) and enrolled after providing written informed consent. Inclusion criteria: males and females, ages 18-60 years, and fluent in written and spoken English. Healthy controls had no history of HS and no first-degree family member with HS. HS patients had a dermatologist confirmed HS diagnosis and documented Hurley Stage of disease severity. Subjects were excluded if they had known allergies to cyanoacrylates, used topical antimicrobials (benzoyl peroxide, chlorhexidine, clindamycin) in the past 2 weeks or systemic antibiotics (oral or IV) in the previous month. Women who were pregnant or breast feeding and individuals with any cognitive impairment were also excluded.

The following information was collected at the time of skin sampling from each subject: 1) sex, 2) age, 3) ethnicity (Hispanic or Latino), 4) Race, 5) alcohol use (frequency: daily, weekly, casual use), 6) tobacco use (current or past smoker or smokeless tobacco; start date; packs/day), 7) personal hygiene products (use of soaps and deodorants, frequency, and brand (if known)) and 8) past and current medications (medication, dose, route, frequency, start/end date).

Additionally, we collected the following information from all HS subjects regarding their history of HS: 1) Date of onset, 2) involved body areas, 3) family history of HS (yes or no; relationship), 4) patient or family history of dissecting cellulitis of the scalp, acne conglobata, pilonidal cysts, pyoderma gangrenosum, or severe acne (If yes, is the condition currently active), 4) any past or current HS specific medications, including biologics (medication, dose, route, frequency, start/end date), and Hurley Stage of disease.

Subject demographics are shown in **Table S1**.

Sampling Method: Cyanoacrylate glue

Sampling occurred within the Dermatology Clinical Research Unit at Penn State Health. Cyanoacrylate glue (SuperGlue, Loctite®, Henkel Corporation, Rocky Hill, CT, USA) follicular biopsy was performed based on methods described (Hall et al. 2018; Mills and Kligman 1983). Fifteen microliters of glue was placed on sterile glass slide, using sterile filter pipette tips, and allowed to dry for ~15 seconds. The glass slide was then pressed onto the subjects' intact skin and held in place for ~1 minute resulting in the glue spreading to ~3 $cm^2$. The glass slide and glue were slowly removed from the axilla or groin by holding the volunteer's skin taut during removal. Lesional skin was an intact red/inflamed nodule and non-lesional skin was at least 5cm away from the lesion and appeared clinically normal. Slides were placed in sterile 50mL conical tube at room temperature.

Using a dissection microscope, follicular "casts" were identified and removed from the superglue using sterile forceps. Casts were counted, placed in a yeast lysis buffer (Epicentre, Illumina), within a sterile 1.5mL microcentrifuge tube and stored at –80°C. For negative control sample, fifteen microliters of glue was placed on sterile glass slide, using sterile filter pipette tips, placed "facing up" on a sterile gauze pad on the exam room counter, left to dry (~3 minutes), and placed in a sterile 50 mL conical tube at room temperature. To simulate "cast collection" on the negative control sample, sterile forceps were touched to the dried glue's surface and the tips of the forceps were dipped into yeast lysis buffer within the collection tube. As with the subject samples, negative control sample was stored at -80C until further processing.

<u>Sampling Method: Swab</u>
Sampling occurred within the Dermatology Clinical Research Unit at Penn State Health.
Sterile synthetic applicators (with plastic handle) were pre-moistened with yeast lysis buffer (Epicentre, Illumina). A 6 $cm^2$ area in the axilla and groin were scrubbed with the swab for thirty seconds and placed in a sterile 1.5 mL tube containing 600 µL of yeast lysis buffer (Epicentre, Illumina) within a sterile 1.5mL microcentrifuge tube. The swabs were immediately stored at -80°C. Lesional skin was an intact red/inflamed nodule and non-lesional skin was at least 5cm away from the lesion and appeared clinically normal.

For negative control sample, a sterile swab was removed from its packaging, held in the air for ~30 seconds to mimic the time that a swab is in contact with human skin, and placed into a sterile 1.5mL tube containing 600 µL of yeast lysis buffer and stored at -80C with all samples.

<u>Controls</u>
Positive mock community control was purchased from ATCC (cat #MSA-1002) and processed. Four negative controls were collected during sample processing.

**Supplemental Methods M2. DNA extraction.**

All samples, including negative control samples, stored at -80°C were removed, thawed on ice, subjected to 0.5 mg lysozyme, and heated to 37°C for 30 mins. Mechanical disruption was performed using a BeadBeater (BioSpec Products) with 0.1 mm sterile glass beads for 3 mins followed by a proteinase K incubation for 30 mins at 65°C. Finally, proteins were removed from samples with MPC reagent (Epicentre, Illumina) and cleared using centrifugation. Supernatants were collected and 350 µL of 70% ethanol was added. DNA was column purified following the DNeasy blood/tissue kit protocols according to manufacturer's instructions (Qiagen). DNA samples were stored at -80°C until sequencing. DNA concentration of each sample was determined by High Sensitivity dsDNA assay by Qubit Fluorometer 3.0 (ThermoFisher). DNA concentrations for negative sampling controls were consistently "below the limit of detection" for this DNA quantification assay.

**Supplemental Methods M3. 16S rRNA gene V3-V4 Sequencing**

The sequences of primers used for the first amplicon PCR are as follows. Both were prepared by IDT DNA Technologies (Coralville, IA, USA):

***PCR Forward Primer (50 bp)***
5'–TCGTCGGCAGCGTCAGATGTGTATAAGAGACAGCCTACGGGNGGCWGCAG–3'
***PCR Reverse Primer (55 bp)***
5'–GTCTCGTGGGCTCGGAGATGTGTATAAGAGACAGGACTACHVGGGTATCTAATCC–3'

16S rRNA gene sequencing was performed at the Genome Sciences and Bioinformatics Core of the Institute of Personalized Medicine at the Penn State College of Medicine. Qubit (Life Technologies) quantified genomic DNA (up to 12.5 ng each) was subjected to amplify V3 and V4 variable regions of the 16S rRNA gene using a two-step, tailed PCR approach that has been validated by Illumina (Illumina, Inc.). Desired size/amount of the amplicon (~550 base pairs (bp)) was confirmed by a Bioanalyzer High Sensitivity DNA chip (Agilent Technologies, Santa Clara, CA, USA) and PCR products were cleaned up by AMPure XP beads (Beckman

Coulter, Jersey City, NJ, USA). If amplicon amount was not sufficient, 5-10 more PCR cycles were added, followed by confirmation with Bioanalyzer High Sensitivity DNA chip and AMPure XP bead cleanup. Approximately 0.5 ng of this initial amplicon was subjected to the second PCR using Nextera XT Index Kits (Illumina, Inc.), which utilizes a dual-index strategy involving 12 (i7) and 8 (i5) indexes, for generating a total of 96 (12 × 8) different index combinations. HiFi HotStart ReadyMix was used along with 10% (v/v) each of i7 and i5 adapters and thermal cycling. Desired size/amount of the amplicon (~630 bp) was confirmed by a Bioanalyzer High Sensitivity DNA chip and PCR products were cleaned up by AMPure XP beads. The resulting libraries were further qualified and quantified by Bioanalyzer High Sensitivity DNA chip, Qubit, and Kapa Library Quantification Kit-Illumina (Kapa Biosystems).

Equimolar libraries were pooled and subjected to Illumina MiSeq sequencing at 2x300 bp (v3 chemistry) supplemented with 5% PhiX spike-in libraries (Illumina Inc.). Illumina CASAVA version 1.8 was used to extract de-multiplexed sequencing reads.

**Supplemental Methods M4: Sequence quality control, processing and analyses.**

16S rRNA gene Fastq processing and OTU table generation

**Diagram**

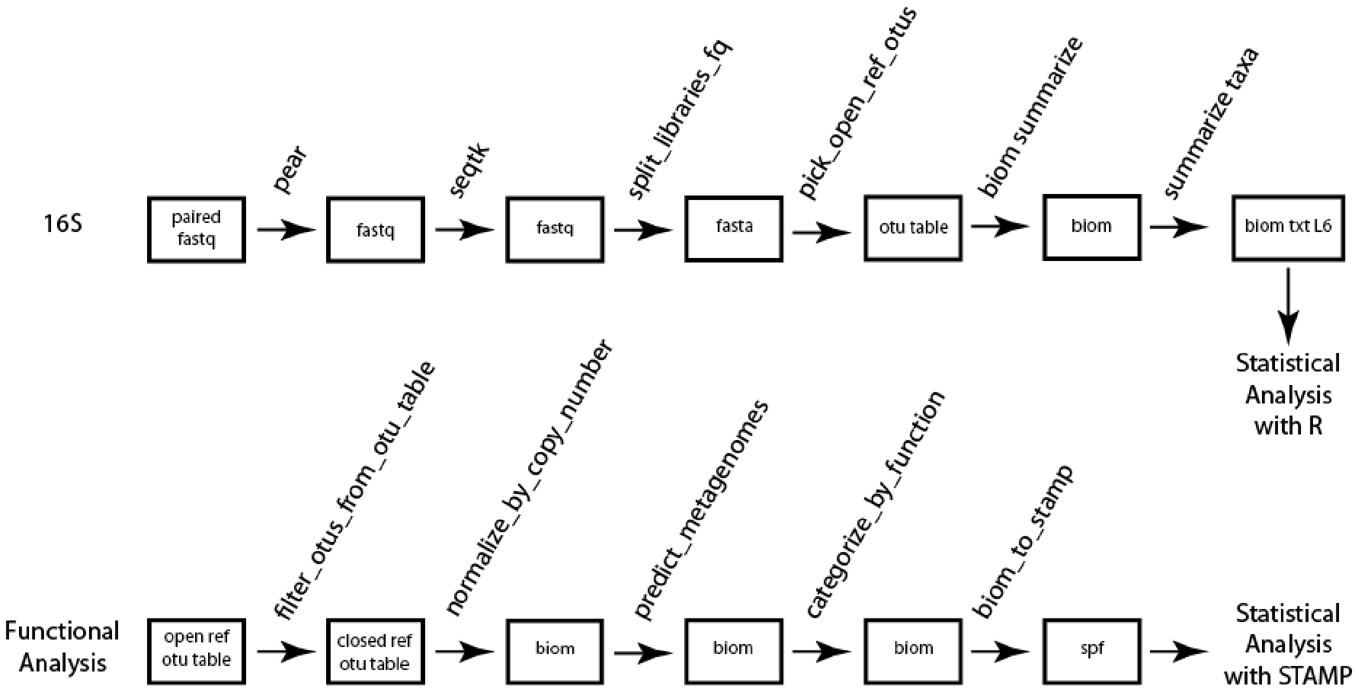

Pear (version 0.9.6; https://sco.h-its.org/exelixis/web/software/pear/) was used to join paired end reads (default base quality PHRED score 33). Each sample was sequenced twice; therefore we merged run 1 and run 2 for each sample to increase reads. Subsampling was then performed at a depth of 75k reads using seqtk (version 1.0-r82; https://github.com/lh3/seqtk). Samples that did not meet this minimum requirement were removed from analysis at this point (**Supplemental Table S2**).

Operational Taxonomic Units (OTUs) were determined using QIIME 1 scripts (version 1.9.1_py2.7.11) (Caporaso et al. 2010), files were converted to fna using 'split_libraries_fastq.py' and then reads were mapped to OTUs with ‘pick_open_reference_otus.py’ with GreenGenes 16S rRNA gene reference database (version 13.8) at 97% sequence similarity (McDonald et al. 2012) and default OTU picking method uclust (Edgar 2010).

Data was converted to a biom format (http://biom-format.org/) to the genus level (L6) with ‘summarize_taxa.py’. All files were merged using ‘merge_otu_tables.py’ and converted to a text file for statistical analyses in R.

Controls

Positive mock community control was (purchased from ATCC MSA-1002) processed as samples above.

Four Negative controls were collected during sample processing. Samples underwent Pear and run 1 and 2 combined as above. Any OTU that did not have 0.5% relative abundance in at least one sample was removed. Genera that hit an average of greater than 2% abundance in all 4 negative controls were removed from processing following the methods described by (Flores et al. 2012), This includes the removal of *Enterococcus*, *Actinomyces*, *Bacillus*, *Streptococcus*, *Sphingomonas*, and *Salinivibrio*. While the average of *Prevotella*, *Anaerococcus*, *Peptoniphilus*, and *Dialister* was > 2% we included these genera because 3 of 4 negative controls were below 2% and one control sample was a clear statistical outlier (determined by 1.5*IQR, bold in table) affecting the abundance.

| | NC 1 (%) | NC 2 (%) | NC 3 (%) | NC 4 (%) | Average (%) |
|---|---|---|---|---|---|
| Prevotella | **32.209** | 0.797 | 0.331 | 0.000 | 8.334 |
| Anaerococcus | **8.774** | 0.018 | 0.166 | 0.008 | 2.242 |
| Peptoniphilus | **9.066** | 0.009 | 0.546 | 0.017 | 2.409 |
| Dialister | **8.839** | 0.000 | 0.007 | 0.000 | 2.212 |
| Enterococcus | 0.854 | 12.207 | 0.273 | 15.760 | 7.273 |
| Actinomyces | 0.634 | 6.957 | 0.331 | 8.197 | 4.030 |
| Bacillus | 0.253 | 7.602 | 1.275 | 4.675 | 3.451 |
| Streptococcus | 0.834 | 10.283 | 1.501 | 7.804 | 5.106 |
| Sphingomonas | 3.583 | 0.049 | 81.794 | 0.004 | 21.357 |
| Salinivibrio | 0.847 | 8.238 | 0.134 | 10.946 | 5.041 |

Functional Analyses

Functional differences of OTUs was determined using PICRUSt (Langille et al. 2013). The OTU table generated in QIIME was first filtered to include only closed reference OTUs at 97% similarity (Greengenes version 13.8). PICRUSt scripts normalize_by_copy_number.py, predict_metagenomes.py, and categorize_by_function.py were used to generate functional data to level 3 functional categories of the Kyoto Encyclopedia of Genes and Genomes (KEGG) orthologs (Kanehisa and Goto 2000). Statistical Analysis of Metagenomic Profile (STAMP) package (Parks et al. 2014) was used to make comparisons between multiple groups Normal, HSN, and HSL using Kruskal-Wallis H-test with the Bonferonni correction. The Microbiome Helper (Comeau et al. 2017) provided the biom_to_stamp.py script used in data processing. The nearest sequenced taxon index (NSTI) was calculated with predict_metagenomes.py to assess quality of our comparisons, all scores were less than 0.15 and all but 7 samples were less than 0.10 indicating good quality comparisons.

| #Sample | Weighted NSTI | #Sample | Weighted NSTI | #Sample | Weighted NSTI | #Sample | Weighted NSTI |
|---|---|---|---|---|---|---|---|
| N001AG | 0.0431 | N008GS | 0.0427 | HS004LAG | 0.0389 | HS007NGS | 0.0378 |
| N001AS | 0.0421 | N009AG | 0.0356 | HS004LAS | 0.0379 | HS008LAG | 0.0427 |
| N001GG | 0.0901 | N009AS | 0.0462 | HS004LGG | 0.0619 | HS008LAS | 0.0537 |
| N001GS | 0.0518 | N009GG | 0.0411 | HS004NAG | 0.0425 | HS008LGG | 0.1002 |
| N002AS | 0.0344 | N009GS | 0.0382 | HS004NAS | 0.0372 | HS008LGS | 0.0490 |
| N002GG | 0.1392 | N010AS | 0.0410 | HS004NGG | 0.0550 | HS008NAG | 0.0450 |
| N002GS | 0.0635 | N010GS | 0.0405 | HS005LAG | 0.0557 | HS008NAS | 0.0300 |
| N003AG | 0.0426 | HS001LAG | 0.0328 | HS005LAS | 0.0994 | HS008NGG | 0.0603 |
| N003AS | 0.0424 | HS001LAS | 0.0326 | HS005LGG | 0.0598 | HS008NGS | 0.0759 |
| N003GG | 0.0835 | HS001LGG | 0.0402 | HS005LGS | 0.0374 | HS009LAS | 0.0390 |
| N003GS | 0.0484 | HS001NAG | 0.0396 | HS005NAG | 0.0867 | HS009LGS | 0.0454 |
| N004AG | 0.0380 | HS001NAS | 0.0343 | HS005NAS | 0.1053 | HS009NAS | 0.0508 |
| N004AS | 0.0389 | HS001NGG | 0.0423 | HS005NGG | 0.1486 | HS009NGS | 0.0435 |
| N004GS | 0.0454 | HS002LAS | 0.0566 | HS005NGS | 0.0363 | HS010LAG | 0.0708 |
| N005AG | 0.0523 | HS002LGG | 0.0331 | HS006LAG | 0.0610 | HS010LGG | 0.0792 |
| N005AS | 0.0795 | HS002LGS | 0.0473 | HS006LAS | 0.0640 | HS010LGS | 0.0844 |
| N005GG | 0.0650 | HS002NAS | 0.0618 | HS006LGS | 0.1251 | HS010NAG | 0.0809 |
| N005GS | 0.0772 | HS002NGG | 0.0312 | HS006NAG | 0.0472 | HS010NGG | 0.0937 |
| N006AG | 0.0457 | HS002NGS | 0.0295 | HS006NAS | 0.0601 | HS010NGS | 0.0606 |
| N006AS | 0.0401 | HS003LAG | 0.0544 | HS006NGS | 0.0753 | HS011LAG | 0.0504 |
| N006GG | 0.0454 | HS003LAS | 0.0451 | HS007LAG | 0.0524 | HS011LGG | 0.0630 |
| N006GS | 0.0394 | HS003LGG | 0.0539 | HS007LAS | 0.0200 | HS011LGS | 0.0720 |
| N007AS | 0.0501 | HS003LGS | 0.0515 | HS007LGG | 0.0938 | HS011NAG | 0.1025 |
| N007GG | 0.0969 | HS003NAG | 0.0530 | HS007LGS | 0.0322 | HS011NGG | 0.0714 |
| N007GS | 0.0502 | HS003NAS | 0.0439 | HS007NAG | 0.0656 | HS011NGS | 0.0742 |
| N008AG | 0.0378 | HS003NGG | 0.0633 | HS007NAS | 0.0340 | | |
| N008AS | 0.0720 | HS003NGS | 0.0571 | HS007NGG | 0.1290 | | |

AG - Axilla Glue; AS - Axilla Swab; GG - Groin Glue; GS - Groin Swab; NAG -Non-Lesion Axilla Glue; NAS - Non-Lesion Axilla Swab; NGG - Non-Lesion Groin Glue; NGS - Non-Lesion Groin Swab; LAG - Lesion Axilla Glue; LAS - Lesion Axilla Swab; LGG - Lesion Groin Glue; LGS - Lesion Groin Swab

#### Supplemental References (for Supplementary Materials)

Caporaso JG, Kuczynski J, Stombaugh J, Bittinger K, Bushman FD, Costello EK, et al. QIIME allows analysis of high-throughput community sequencing data. Nat. Methods. NIH Public Access; 2010;7(5):335–6

Comeau AM, Douglas GM, Langille MGI. Microbiome Helper: a Custom and Streamlined Workflow for Microbiome Research. mSystems. American Society for Microbiology Journals; 2017;2(1):e00127-16

Edgar RC. Search and clustering orders of magnitude faster than BLAST. Bioinformatics. Oxford University Press; 2010;26(19):2460–1

Flores GE, Henley JB, Fierer N. A Direct PCR Approach to Accelerate Analyses of Human-Associated Microbial Communities. Gilbert JA, editor. PLoS One. Public Library of Science; 2012;7(9):e44563

Hall JB, Cong Z, Imamura-Kawasawa Y, Kidd BA, Dudley JT, Thiboutot DM, et al. Isolation and Identification of the Follicular Microbiome: Implications for Acne Research. J. Invest. Dermatol. Elsevier; 2018;138(9):2033–40

Kanehisa M, Goto S. KEGG: kyoto encyclopedia of genes and genomes. Nucleic Acids Res. Oxford University Press; 2000;28(1):27–30

Langille MGI, Zaneveld J, Caporaso JG, McDonald D, Knights D, Reyes JA, et al. Predictive functional profiling of microbial communities using 16S rRNA marker gene sequences. Nat. Biotechnol. Nature Publishing Group; 2013;31(9):814–21

McDonald D, Price MN, Goodrich J, Nawrocki EP, DeSantis TZ, Probst A, et al. An improved Greengenes taxonomy with explicit ranks for ecological and evolutionary analyses of bacteria and archaea. ISME J. Nature Publishing Group; 2012;6(3):610–8

Mills OH, Kligman AM. The follicular biopsy. Dermatologica. 1983;167(2):57–63

Parks DH, Tyson GW, Hugenholtz P, Beiko RG. STAMP: statistical analysis of taxonomic and functional profiles. Bioinformatics. Oxford University Press; 2014;30(21):3123–4
